# Kv2 conductances are not required for C-bouton mediated enhancement of motoneuron output

**DOI:** 10.1101/2022.07.23.501232

**Authors:** Calvin C. Smith, Robert M. Brownstone

## Abstract

Neural motor systems have evolved complex circuits that afford animals a range of behaviours essential for survival. C-bouton synapses arising from cholinergic V0_C_ interneurons amplify motoneuron activity via muscarine type 2 receptors, thus increasing muscle contraction force. Recent work in neonatal mouse motoneurons suggests that delayed rectifier currents carried by post-synaptically clustered K_V_2.1 channels are crucial to C-bouton amplification. Here we use a motoneuron conditional K_V_2.1 knockout to show that while K_V_2.1 modulates maximal firing in neonatal mice, its removal minimally affects either mature motoneuron firing or the enhanced firing rates in response to exogenously applied muscarine. In keeping with this, pharmacological blockade of K_V_2 currents has minimal electrophysiological effects on mature motoneurons. Furthermore, amplification of electromyography activity during high force tasks was unchanged following K_V_2.1 deletion. We next show that K_V_2.2 is also expressed by spinal motoneurons and colocalises with K_V_2.1 opposite C-boutons. We suggest that the primary function of K_V_2 proteins – K_V_2.1 and K_V_2.2 – is non-conducting in motoneurons, and that K_V_2.2 can function in the absence of K_V_2.1, perhaps to ensure the integrity of the synapse.

## Introduction

For most mammals, survival is dependent upon the ability to produce a broad spectrum of adaptable behaviours. These behaviours are produced by spatiotemporal patterns of muscle contractions, which are governed by the motoneurons that innervate them. Thus, motoneuron activity must be regulated so as to support such diverse motor outputs. In addition to the properties of the muscle fibre types, the strength of muscle contraction is governed by the number of active innervating motoneurons and their firing frequencies (Enoka & Farina, 2021). Thus, to understand the neural mechanisms of behaviour, it is essential to understand how motoneurons produce repetitive spike trains.

The relationship between the inputs a motoneuron receives and its outputs is often represented by a “frequency-current” or ƒ-I curve, with the current being that injected through a recording electrode. The slope(s) of the linear segment(s) (gain) of this curve can be altered in a task-dependent manner by neuromodulator systems. One system that can increase the gain of motoneurons is that comprised of presynaptic C-boutons, named because of their association with specialised endoplasmic reticulum (ER) called “subsurface cisterns” (Conradi, 1969), and which are terminals of cholinergic V0c interneurons (Miles *et al*., 2007; Zagoraiou *et al*., 2009). The postsynaptic motoneuron membrane apposing C-boutons includes clusters of many proteins, including type 2 muscarinic acetylcholine (M2) receptors (Hellström *et al*., 2003), slow calcium-dependent potassium channels (SK2, SK3; Deardorff *et al*., 2013), and voltage-gated delayed rectifier potassium channels (Kv2.1; Muennich & Fyffe, 2004). These synapses mature over the first three weeks of post-natal development in mice in parallel with the development of weight-bearing locomotor function (Wilson *et al*., 2004). And in the adult mouse, this system is recruited for high force outputs, such as the extensor stroke in swimming (Zagoraiou *et al*., 2009). But how activation of M2 receptors leads to the increase in excitability needed for these tasks remains elusive, leaving a significant hole in our understanding of movement.

It has been hypothesised that M2 receptor activation affects K_V_2.1 function to actuate C-bouton-mediated amplification (Muennich & Fyffe, 2004; Romer *et al*., 2014; Fletcher *et al*., 2017; Romer *et al*., 2019; Nascimento *et al*., 2020; Deardorff *et al*., 2021). K_V_2 channels are widely expressed through the central nervous system (Trimmer, 1991). There are two main K_V_2 subunits, KCNB1 (K_V_2.1) and KCNB2 (K_V_2.2), which share similar biophysical properties, and are often, but not always, co-expressed (Frech *et al*., 1989; Hwang *et al*., 1992; Hwang *et al*., 1993a; Hwang *et al*., 1993b). As delayed rectifiers, K_V_2 channels are crucial to regulating neuronal excitability in some neurons. Their importance in neuronal function is evident by clinical reports of people with KCNB1 mutations, who have a myriad of problems including reduced cognitive capacity and epilepsy (Torkamani *et al*., 2014; Thiffault *et al*., 2015; Miao *et al*., 2017). About one-half of the 26 reported people are hypotonic, and 2/3 of those were reported in early life. On the other hand, mice with KCNB1 deletions are hyperactive: their problems are not in motoneuron function per se, and they are not hypotonic (Speca *et al*., 2014). These results do not support the hypothesis that K_V_2.1 channels are required for motor output. And they do not shed light on what the role of these channels in motoneurons might be.

Using K_V_2 channel blockers (K_V_2.1 and K_V_2.2), several groups have suggested significant roles for K_V_2.1 conductances in C-bouton-mediated amplification in neonatal rodent motoneurons (Fletcher *et al*., 2017; Romer *et al*., 2019; Nascimento *et al*., 2020). Although results were not all entirely consistent between the studies, overall they suggested that when active, C-boutons recruit local K_V_2.1 channels to maintain narrow spikes and fast afterhyperpolarisation (fAHP) amplitudes, thus supporting high frequency firing by preventing Na^+^ channel inactivation and the resultant depolarisation block. That is, current work suggests K_V_2.1 channels have significant conducting roles.

Non-conducting roles have also been proposed for K_V_2.1 channels (Johnson *et al*., 2018; Kirmiz *et al*., 2018; Deardorff *et al*., 2021), as they form physical links between the plasma membrane (PM) and ER by binding to VAP (VAMP associated protein) proteins (Fox *et al*., 2015; Johnson *et al*., 2018; Kirmiz *et al*., 2018). Given the calcium-dependence of proteins clustered at the C-bouton, and that subsurface cisterns serve as regulators of Ca^+^ micro domains, K_V_2.1 could provide a link allowing M2 receptors to modulate local Ca^2+^ concentration and therefore channel function. PM-ER junctions have not been explicitly investigated in motoneurons, but Deardorff *et al*. (2021) recently provided evidence for VAP expression in this post-synaptic domain.

Taken together, the role of motoneuron K_V_2.1 channels in regulating motoneuron firing and in C-bouton function remains unclear, at least in part due to the paucity of experiments assessing channel function in mature motoneurons and in awake behaving animals.

In this study, our grandiose aim was to define K_V_2.1 function in mature motoneurons from electrophysiology through to animal behaviour. We used a conditional knock-out (cKO) mouse in which cholinergic neurons (including motoneurons) lacked K_V_2.1 channels. We then used whole cell patch clamp electrophysiology to compare the firing characteristics of mature K_V_2.1^ON^ (control) and K_V_2.1^OFF^ (ChAT-K_V_2.1^OFF^) motoneurons and the effects of the specific K_V_2 channel blocker Guangxitoxin-1E (GxTX-1E) on these properties. We repeated these experiments in early post-natal control motoneurons to assess whether developmental clustering of K_V_2.1 channels influences their role in regulation of firing. To determine the role of K_V_2.1 in C-bouton function, we activated M2 receptors *in vitro* using muscarine and compared excitability changes in control and K_V_2.1 cKO motoneurons. Finally, to assess whether K_V_2.1 cKO influences motor amplification, we studied high force output behaviours whilst recording electromyography (EMG) activity in hind limb muscles. We found surprisingly little effect of K_V_2.1 conductances on motoneuron physiology or behaviour and suggest that K_V_2 channels play a primarily non-conducting role. Furthermore, we suggest that in the absence of K_V_2.1, K_V_2.2 can subsume the non-conducting roles of K_V_2 channels including maintaining the PM-ER relationship.

## Results

### Conditional knockout of K_V_2.1

In order to investigate the contribution of K_V_2.1 channels to motoneuron physiology, we aimed to eliminate K_V_2.1 from motoneurons through multi-generational crossing of ChAT-IRES-Cre (ChAT^(Cre/Cre)^) mice with homozygous floxed *KCNB1* mice (*KCNB1*^(f/f)^) to generate ChAT^(Cre/wt)^;*KCNB1*^(-/-)^, ChAT-K_V_2.1^OFF^ mice (Figure 1-figure supplement 1). To test whether the strategy effectively eliminated K_V_2.1 channels from cholinergic neurons, we proceeded with immunohistochemical labelling using antibodies against ChAT and K_V_2.1 in ChAT-K_V_2.1^OFF^ and littermate controls (ChAT^(wt/wt)^;*KCNB1*^(f/f)^, Figure 1A-E2). Our analyses showed similar K_V_2.1 punctae density in the dorsal horn of both genotypes (mean difference = -0.04 /100 µm^2^, Hedges g = - 0.19, CI_95_ = [-1.16, 0.76], N = 9, Figure 1F), but large reductions in the intermediate (mean difference = -0.11 /100 µm^2^, Hedges g = -1.47, CI_95_ = [-2.25, -0.58], N = 9, Figure 1G) and ventral horns (mean difference = -0.15 /100 µm^2^, Hedges g = -1.87, CI_95_ = [-2.50, -1.22], N = 9, Figure 1H). This result validates the strategy, given that cholinergic neurons are found in both these regions, most densely in the ventral horn where motoneurons reside. We found no K_V_2.1 labelling on motoneuron somata of ChAT-K_V_2.1^OFF^ mice, suggesting that any residual punctae in the ventral horns were associated with non-cholinergic neurons (Figure 1D-E2). Thus, this strategy for removing K_V_2.1 channels from motoneurons was successful.

**Figure 1.**
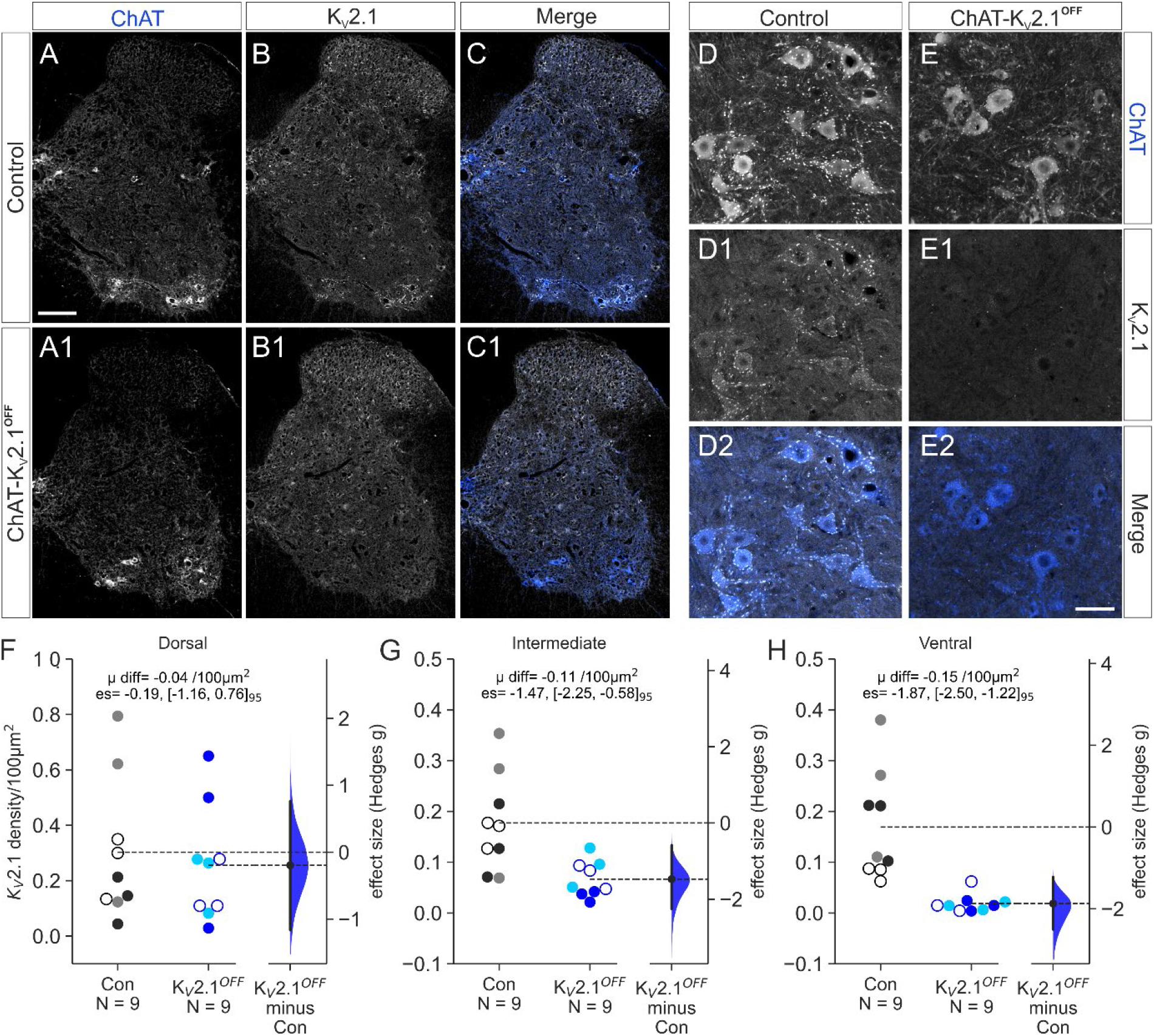
Conditional knockout of K_V_2.1 in ChAT^+^ lumbar motoneurons. Confocal photomicrographs (20x tile scans, 1μm optical section) of spinal cord hemisections from a control (A-C) and ChAT-K_V_2.1**^OFF^** mouse **(A1-C1)**. **(D-D2)** 20x z-stack images from motor pool of a control mouse spinal cord. **(E-E2)** As in D-D2, but from a ChAT-K_V_2.1**^OFF^**mouse. Note the absence of K_V_2.1^+^ puncta on motoneurons following conditional knockout**. (F-H)** Gardner-Altman estimation plots showing K_V_2.1 density in dorsal (**F**), intermediate (**G**) and ventral (**H**) regions. Density was calculated by counting the number of K_V_2.1 punctae within a region of interest (ROI). The dimensions of the ROI were the same for each section. In each Gardner-Altman plot, Control (Con) and ChAT-K_V_2.1**^OFF^** groups are plotted on the left axes and the bootstrapped sampling distribution (5000 reshuffles) for Hedges g effect size are plotted on the right. Each point represents one section, with different fill colours representing sections from different animals. The Hedges g effect sizes are depicted as a dot; the 95% confidence interval is indicated by the ends of the vertical error bar. The mean difference (μ diff), effect size (es) and 95% confidence intervals [lower, upper] are displayed at the top of each plot. Scale bar = 100 μm in A-C1 and 40 μm in D-F1. Three random, free floating slices (30 μm thickness) were quantified per animal (Control, N=9 slices from 3 animals; ChAT-KV2.1**^OFF^**, N=9 slices from 3 animals, all females).

### Firing characteristics are similar in control and K_V_2.1^OFF^ motoneurons

Having confirmed that ChAT-K_V_2.1^OFF^ motoneurons are devoid of K_V_2.1, we next performed whole cell current clamp experiments to determine whether ɑ-motoneuron firing characteristics were affected by channel absence.

We first assessed the passive membrane properties of large control and K_V_2.1^OFF^ motoneurons, finding a small mean difference in **resting membrane potential** (RMP, K_V_2.1^OFF^ = -68 ± 3 mV, N = 29; control = -67 ± 3 mV, N = 36, mean difference = -1 mV, Hedges g = -0.49, CI_95_ = [-0.91, 0.01], Figure 2 - figure supplement 1A), while **Resistance** (K_V_2.1^OFF^ = 21 ± 11 MΩ, N = 28; control = 25 ± 12 MΩ, N = 35, mean difference = -4 MΩ, Hedges g = -0.31, CI_95_ = [-0.79, 0.18]; Figure 2 - figure supplement 1B), **Capacitance** (K_V_2.1^OFF^ = 522 ± 326 pF, N = 28; control = 429 ± 160 pF, N = 35, mean difference = 93 pF, Hedges g = 0.36, CI_95_ = [-0.10, 0.76]; Figure 2 - figure supplement 1C), **Time constant** (ChAT-K_V_2.1^OFF^ = 9.2 ± 3.4 ms, N = 28; control = 9.3 ± 2.7 ms, N = 35; mean difference = -0.1 ms, Hedges g = -0.02, CI_95_ = [-0.54, 0.50]; Figure 2 - figure supplement 1D), and **Rheobase** (K_V_2.1^OFF^ = 1.2 ± 0.8 nA, N = 28; control = 1.1 ± 0.7 nA, N = 35, mean difference = 0.03 nA, Hedges g = 0.05, CI_95_ = [-0.44, 0.54]; Figure 2 - figure supplement 1E) were all similar.

Next, we compared the firing capabilities of motoneurons from both groups. The maximum instantaneous frequency of repetitive firing during the entire 500ms pulse (**Overall frequency**, Figure 2A-C, A1-B1) was higher (mean difference = 17 Hz) in K_V_2.1^OFF^ motoneurons (87 ± 23 Hz, N = 29) compared to control (70 ± 23 Hz, N = 36, Hedges g = 0.72, CI_95_ = [0.18, 1.2], Figure 2C), however the slope of the ƒ-I curve was similar between groups (**Overall frequency slope**, mean difference = 2 Hz/nA, K_V_2.1^OFF^ = 24 ± 14, N = 29; control = 22 ± 10 Hz/nA, N = 36, Hedges g = 0.14, CI_95_ = [-0.33, 0.63], Figure 2C1). Taken together, these results suggest that while there is no difference in excitability of the two populations of motoneurons, K_V_2.1 conductances may limit maximum firing rates. The initial instantaneous frequencies (of first two spikes) were similar (**1^st^ interval frequency**, K_V_2.1^OFF^ = 210 ± 49 Hz, N = 29; control = 205 ± 42 Hz, N = 36, mean difference = 6 Hz, Hedges g = 0.12, CI_95_ = [-0.38, 0.62], Figure 2D) as were ƒ-I slopes of this interval (**1^st^ interval slope**, K_V_2.1^OFF^ = 67 ± 31 Hz/nA, N = 29; control = 65 ± 45 Hz/nA, N = 36, mean difference = 2 Hz/nA, Hedges g = 0.04, CI_95_ = [-0.38, 0.61], Figure 2D1).

**Figure 2.**
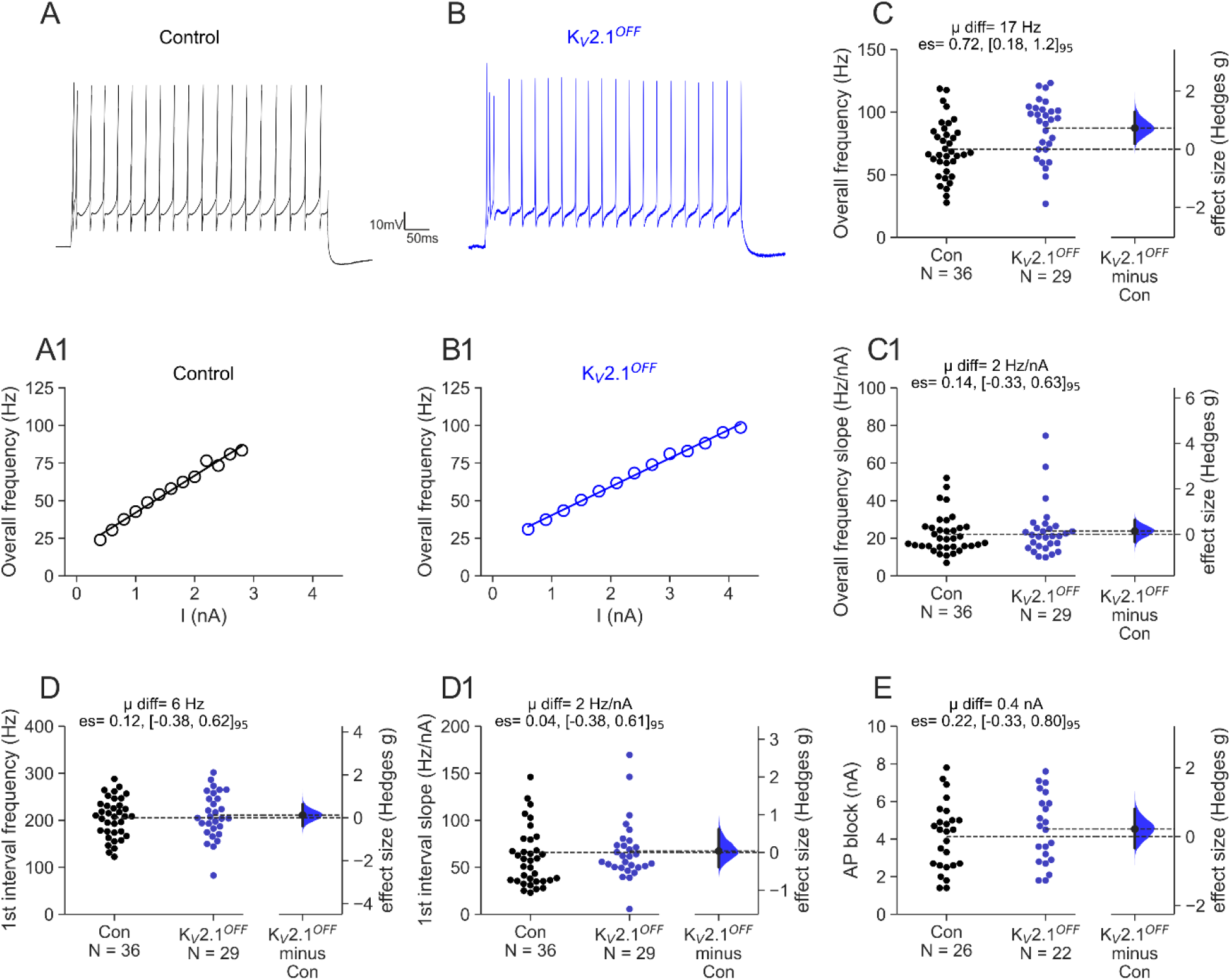
Comparing firing capabilities of ChAT-K_V_2.1^OFF^ with control motoneurons. Patch clamp electrophysiology was used to assess various firing characteristics of lumbar motoneurons in control and ChAT-K_V_2.1**^OFF^** mice. **(A-B)** Representative traces from a control (Con, black) and ChAT-K_V_2.1**^OFF^** motoneuron (blue) at 3 x threshold for repetitive firing. **(A1-B1)** Representative ƒ-I plots for control (black) and ChAT-K_V_2.1**^OFF^**motoneurons (blue). **(C)** Gardner-Altman estimation plots for overall frequency, calculated as the mean instantaneous frequency between all spikes within a 500 ms train. Control and ChAT-K_V_2.1**^OFF^**groups are plotted on the left axes and the bootstrapped sampling distribution (5000 reshuffles) for Hedges g effect size are plotted on the right. Each point represents a single motoneuron. The Hedges g effect size is depicted as a dot; the 95% confidence interval is indicated by the ends of the vertical error bar. The mean difference (μ diff), effect size (es) and 95% confidence intervals [lower, upper] are displayed at the top of each plot. Experimental unit (N) = motoneurons recorded from 23 control (n= 9 females, 14 males) and 14 ChAT-K_V_2.1**^OFF^**mice (n= 7 females, 7 males). **(C1)** The slope of the ƒ-I curve for the overall frequency, calculated from the 1^st^ linear portion of the plot, as indicated by lines fitted in A1-B1. **(D)** The maximum instantaneous frequency of the 1^st^ inter-spike interval in a 500 ms train. **(D1)** As in C1, but the ƒ-I curve is only calculated for the first inter-spike interval. **(E)** The current threshold for depolarising block (spike failure).

In some neurons, K_V_2 delayed rectifier currents ensure Na^+^ channel recovery by maintaining fast action potential (AP) repolarisation kinetics, thereby preventing depolarising block (Liu & Bean, 2014; Kimm *et al*., 2015). We therefore assessed whether cKO of K_V_2.1 affected the current threshold at which membrane depolarisation blocked spike production (Figure 2E). Despite a lack of K_V_2.1, K_V_2.1^OFF^ motoneurons entered depolarising block at similar current thresholds as control motoneurons (K_V_2.1^OFF^ = 4.5 ± 1.8 nA, N = 22; control = 4.1 ± 1.8 nA, N = 26; mean difference = 0.4 nA, Hedges g = 0.22, CI_95_ = [-0.33, 0.80]), suggesting K_V_2.1 contributes little to preventing depolarising block in spinal motoneurons.

### cKO of K_V_2.1 does not alter motoneuron action potential characteristics

K_V_2.1 delayed rectifier currents are important for maintaining spike shape in many neurons, with numerous studies showing that their block increases spike width and reduces amplitude of both the fast AHP (fAHP) and the spike itself (Liu & Bean, 2014; Kimm *et al*., 2015; Fletcher *et al*., 2017). Thus, K_V_2.1^OFF^ motoneurons were expected to have greater action potential ½ widths, with smaller fAHP and spike amplitudes. However, we found that K_V_2.1^OFF^ and control motoneurons had similar **spike amplitudes** (K_V_2.1^OFF^ = 81 ± 7 mV, N = 27; control = 80± 4 mV, N = 36, mean difference = 0.8 mV, Hedges g = 0.13, CI_95_ = [-0.41, 0.68], Figure 3A-B), **AP ½ widths** (K_V_2.1^OFF^ = 0.63 ± 0.12 ms, N = 27; control = 0.64 ± 0.13 ms, N = 36, mean difference = -2e-03 ms, Hedges g = -0.02, CI_95_ = [-0.52, 0.49], Figure 3A,C), and **fAHP amplitudes** (K_V_2.1^OFF^ = 14 ± 6 mV, N = 27; control = 14 ± 5 mV, N = 36, mean difference = -0.3 mV, Hedges g = -0.05, CI_95_ = [-0.54, 0.46], Figure 3D).

**Figure 3.**
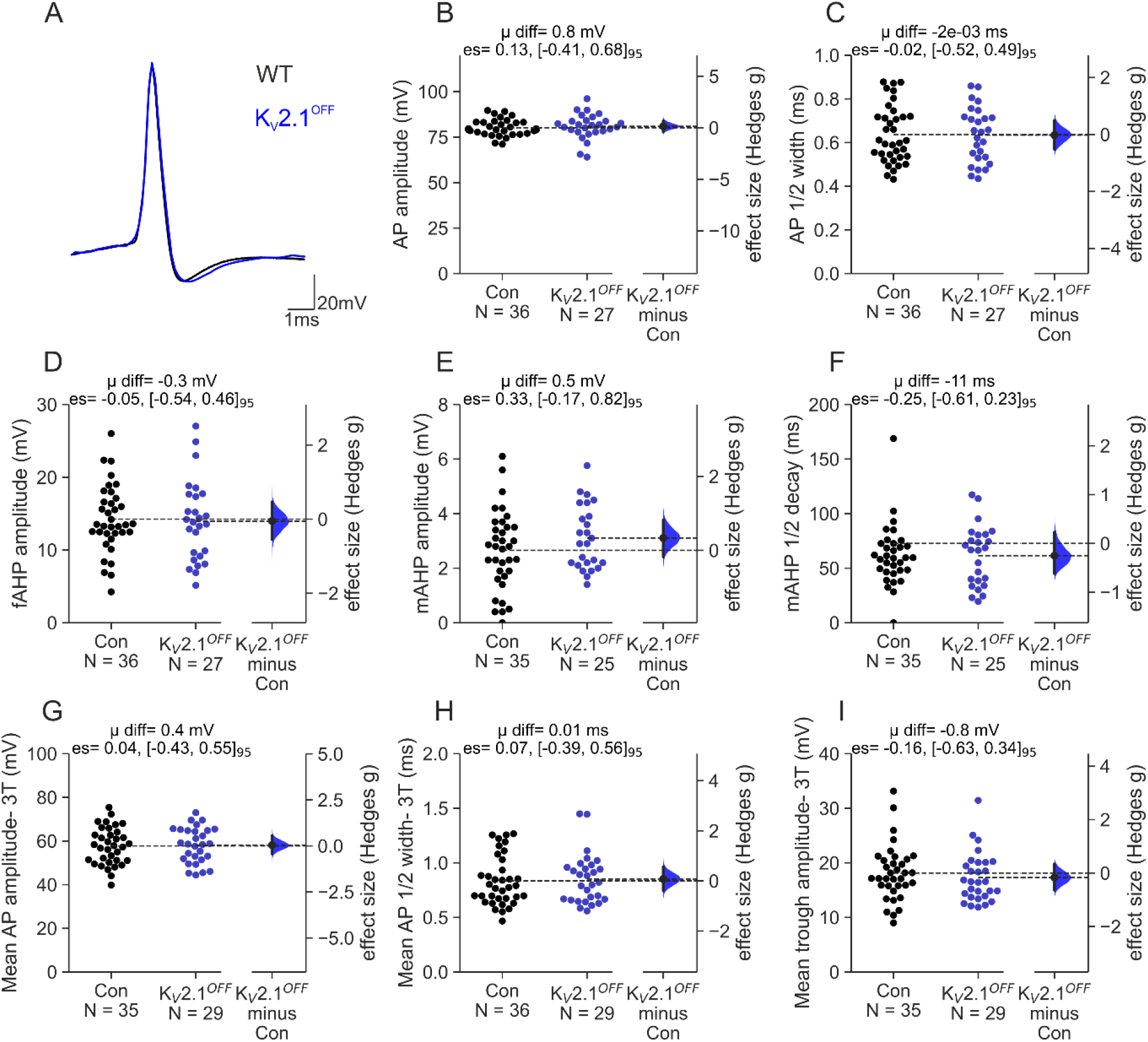
Comparing action potential characteristic of control and ChAT-K_V_2.1^OFF^ motoneurons. **(A)** Representative averages of 15-30 single action potentials (AP) evoked with a 20 ms current pulse in control (Con, black) and ChAT-K_V_2.1**^OFF^** (blue) motoneurons. **(B-F)** Gardner-Altman estimation plots for action potential amplitude **(B)**, ½ width **(C)**, fast afterhyperpolarisation amplitude **(fAHP, D)**, medium afterhyperpolarisation amplitude **(mAHP, E)**, mAHP ½ decay time **(F)**. Control and ChAT-K_V_2.1**^OFF^** groups are plotted on the left axes and the bootstrapped sampling distribution (5000 reshuffles) for Hedges g effect size are plotted on the right. The Hedges g effect sizes are depicted as black dots; the 95% confidence intervals are indicated by the ends of the vertical error bars. The mean difference (μ diff), effect sizes (es) and 95% confidence intervals [lower, upper] are displayed at the top of each plot. **(G-I)** As is B-F, but mean of all APs in a 500ms pulse are taken at 3x threshold (3T) for repetitive firing. AP amplitude is shown in **(G)**, mean AP ½ width-3T **(H)**, and mean AHP-3T **(I)**. Experimental unit (N) = motoneurons recorded from 23 control (n= 9 females, 14 males) and 14 ChAT-K_V_2.1**^OFF^**mice (n= 7 females, 7 males)

Although small conductance calcium activated potassium (SK) currents are the main conductances contributing to the medium AHP (mAHP), K_V_2.1 delayed rectifier currents have recently been suggested as an auxiliary mAHP conductance in motoneurons (Nascimento *et al*., 2020). Our analyses showed no consistent difference in **mAHP amplitudes** (K_V_2.1^OFF^ = 3.1 ± 1.2 mV, N = 25; control = 2.7 ± 1.4 mV, N = 35, mean difference = 0.5 mV, Hedges g = 0.33, CI_95_ = [-0.17, 0.82], Figure 3E) or **½ decay** (K_V_2.1^OFF^ = 61 ± 27 ms, N = 25; control = 73 ± 53 ms, N = 35; mean difference = -11ms, Hedges g = -0.25, CI_95_ = [-0.61, 0.23], Figure 3F).

Unlike single evoked spikes, action potentials during repetitive firing are subject to conductances with longer time constants (Miles *et al*., 2005), and therefore their morphology may be differentially affected (Romer *et al*., 2019). Thus, we also assessed mean AP characteristics at 3 times the threshold for repetitive firing, and found that spike amplitude (**Mean AP amplitude-3T**, K_V_2.1^OFF^ = 58 ± 8mV, N = 29; control = 58 ± 8mV, N = 35; mean difference = 0.4 mV, Hedges g = 0.04, CI_95_ = [-0.43, 0.55], Figure 3G), AP ½ width (**Mean AP ½ width-3T**, K_V_2.1^OFF^ = 0.85 ± 0.23 mV, N = 29; control = 0.83 ± 0.22 ms, N = 36; mean difference = 0.01ms, Hedges g = 0.07, CI_95_ = [-0.39, 0.56], Figure 3H), and interspike trough (**Mean trough amplitude-3T**, K_V_2.1^OFF^ = 17 ± 5 mV, N = 29; control = 18 ± 5mV, N = 35; mean difference = -0.8 mV, Hedges g = 0.16, CI_95_ = [-0.63, 0.34], Figure 3I) were similar in control and K_V_2.1^OFF^ motoneurons.

Together, comparisons of firing characteristics and AP morphology from control and K_V_2.1^OFF^ motoneurons indicate that either K_V_2.1 channels have minimal role in regulating firing capabilities or that there were effective compensatory mechanisms following their loss.

### K_V_2 block by GxTX-1E has minimal effect on motoneuron maximal firing characteristics

Given that various mechanisms could mask associated deficits in K_V_2.1^OFF^ motoneurons, we next assessed the role of K_V_2.1 channels in shaping motoneuron firing characteristics pharmacologically. We used the Chinese tarantula toxin Guangxitoxin-1E (GxTX-1E), which at 100 nM potently and selectively inhibits K_V_2 channels by slowing the opening and closing times (Herrington *et al*., 2006; Liu & Bean, 2014; Fletcher *et al*., 2017; Nascimento *et al*., 2020; Newkirk *et al*., 2022). Because K_V_2.1 channels have recently been suggested to play a significant role in regulating motoneuron firing (Fletcher *et al*., 2017; Romer *et al*., 2019; Nascimento *et al*., 2020), we hypothesised GxTX-1E would significantly alter firing characteristics in control but not K_V_2.1^OFF^ motoneurons.

To confirm that in our hands GxTX-1E does not have significant effects on passive membrane properties of wild-type motoneurons (Fletcher *et al*., 2017; Nascimento *et al*., 2020), we first measured these effects in both our control and K_V_2.1^OFF^ motoneurons by measuring voltage responses to sub-threshold and threshold current steps (Figure 4 - figure supplement 1A-E). In line with previous work, there were no effects of the toxin on **Resting membrane potential** in either condition (**K_V_2.1^OFF^**: nACSF = -68 ± 3 mV, GxTX-1E = -69 ± 4 mV, mean difference = -0.2 mV, Hedges g = - 0.04, CI_95_ = [-0.28, 0.19], N = 12; **control**: nACSF = -65 ± 3 mV, GxTX-1E = -65 ± 3 mV, mean difference = -0.02 mV, Hedges g = -0.02, CI_95_ = [-0.32, 0.30], N = 20, Figure 4 S 1B). Similarly, we found no consistent effect of GxTX-1E on **Resistance** in either control (nACSF = 22 ± 10 MΩ, GxTX-1E = 20 ± 9 MΩ, N = 19, mean difference = -3 MΩ, Hedges g = -0.26, CI_95_ = [-0.58, 0.05], Figure 4 SC) or **Capacitance** (nACSF = 429 ± 170 pF, GxTX-1E = 490 ± 208 pF, mean difference = 61 pF, Hedges g = 0.31, CI_95_ = [-0.001, 0.60], N = 19, Figure 4 - figure supplement 1D) or K_V_2.1^OFF^motoneurons (**Resistance**: nACSF = 19 ± 10 MΩ, GxTX-1E = 19 ± 10 MΩ, mean difference = 0.01 MΩ, Hedges g= 8e-04, CI_95_ = [-0.22, 0.19], N = 11, Figure 4 - figure supplement 1C; **Capacitance**: nACSF = 636 ± 481 pF, GxTX-1E = 595 ± 366 pF, mean difference = -41 pF, Hedges g = -0.09, CI_95_ = [-0.26, 0.31], N = 11, Figure 4 figure supplement 1D).

**Figure 4.**
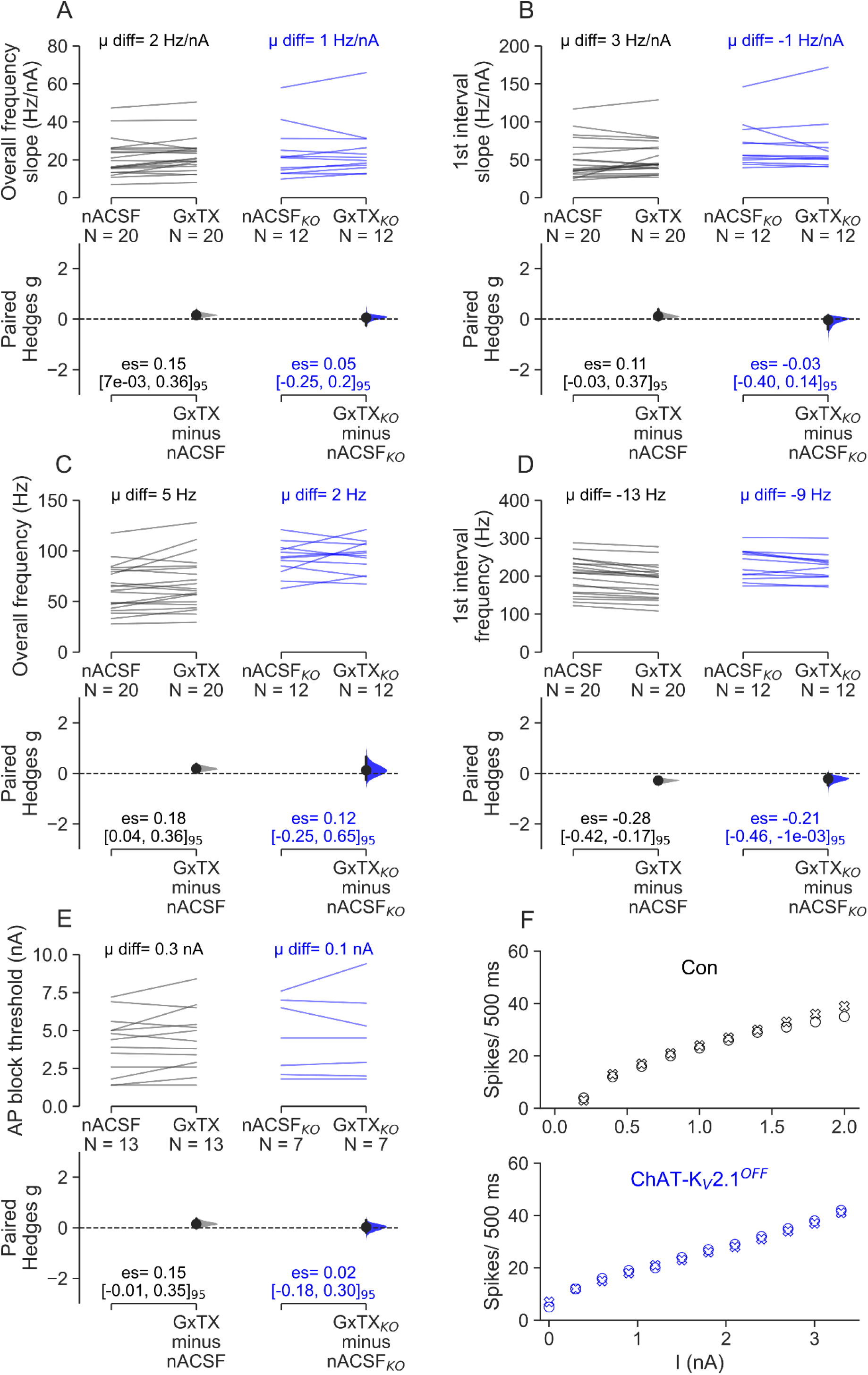
Comparing the effects of GxTX-1E on firing characteristics and excitability of control and ChAT-K_V_2.1^OFF^ motoneurons. **(A-F)** Paired Hedges g for overall frequency slope **(A)**, 1^st^ interval frequency slope **(B)**, maximum overall frequency **(C)**, maximum 1^st^ interval frequency **(D)**, and AP block threshold **(E)** in control (left) and ChAT-K_V_2.1^OFF^ (right) motoneurons are shown in Cumming paired estimation plots. Individual motoneurons are plotted on the upper graphs, with each paired set of observations (nACSF followed by 100 nM GxTX-1E) connected by a line. On the lower plots, effect sizes (Hedges g) are plotted with bootstrapped sampling distribution (5000 reshuffles). Effect sizes are depicted as dots; 95% confidence intervals are indicated by the vertical error bars. The bootstrapped mean differences (μ diff) are shown on the upper plots, and the effect sizes (es) and 95% confidence intervals [lower, upper] are displayed on the lower plots. Experimental unit (N) = motoneurons. **(F)** Scatter plots of spike number in response to increasing current input for representative motoneurons from control (upper graph, black) and ChAT-K_V_2.1**^OFF^** mice (lower graph, blue). Spike number in motoneurons perfused with nACSF only is plotted using ‘O’ and following 10 minutes GxTX-1E is plotted with ‘X’. Number of animals used was as follows: control= 13 (6 females, 7 males), ChAT-K_V_2.1**^OFF^ =** 6 (4 females, 2 males).

In contrast, we saw small increases in the rheobase of both K_V_2.1^OFF^ and control motoneurons, suggesting that GxTX-1E had effects on conductances other than K_V_2.1 (**K_V_2.1^OFF^**: nACSF = 1.3 ± 0.9 nA, GxTX-1E = 1.6 ± 0.9 nA, mean difference = 0.3 nA, Hedges g = 0.29, CI_95_ = [0.11, 0.60], N = 12; **control**: nACSF = 1.3 ± 0.8 nA, GxTX-1E = 1.5 ± 0.9 nA, mean difference = 0.2 nA, Hedges g = 0.24, CI_95_ = [0.09, 0.41], N = 20; Figure 4 - figure supplement 1E).

GxTX-1E block of K_V_2 channels had very little effect on motoneuron repetitive firing properties (Figure 4). The slope of the ƒ-I relationship for overall and initial frequency were unaltered by GxTX-1E in both K_V_2.1^OFF^ (**Overall ƒ-I:** nACSF = 24 ± 14 Hz/nA, GxTX-1E = 25 ± 14 Hz/nA, mean difference = 1 Hz/nA, Hedges g = 0.05, CI_95_ = [-0.25, 0.20], N = 12, Figure 4A, F; **Initial interval ƒ-I**: nACSF = 68 ± 30 Hz/nA, GxTX-1E = 67 ± 37 Hz/nA, mean difference = 1 Hz/nA, Hedges g = - 0.03, CI_95_ = [-0.40, 0.14], N = 12, Figure 4B), and control motoneurons (**Overall ƒ-I:** nACSF = 21 ± 10 Hz/nA, GxTX-1E = 23 ± 10 Hz/nA, mean difference = 2 Hz/nA, Hedges g = 0.15, CI_95_ = [0.01, 0.36], N = 20, Figure 4A; **Initial interval ƒ-I**: nACSF = 49 ± 26 Hz/nA, GxTX-1E = 52 ± 24 Hz/nA, mean difference = 3 Hz/nA, Hedges g = 0.11, CI_95_ = [-0.03, 0.37], N = 20, Figure 4B).

Similarly, we found little effect of GxTX-1E on the peak overall firing frequency for either K_V_2.1^OFF^ (nACSF = 92 ± 16 Hz, GxTX-1E = 95 ± 16 Hz, mean difference = 2 Hz, Hedges g = 0.12, CI_95_ = [- 0.25, 0.65], N = 12, Figure 4C), or control motoneurons (nACSF = 63 ± 23 Hz, GxTX-1E = 68 ± 26 Hz, mean difference = 5 Hz, Hedges g = 0.18, CI_95_ = [0.04, 0.36], N = 20, Figure 4C). However, GxTX-1E did cause relatively small reductions in the firing frequency of the initial spikes for both K_V_2.1^OFF^ (nACSF = 231 ± 40 Hz, GxTX-1E = 222 ± 36 Hz, mean difference = -9 Hz, Hedges g = - 0.21, CI_95_ = [-0.46, -1e-03], N = 12, Figure 4D) and control motoneurons (nACSF = 199 ± 47 Hz, GxTX-1E = 186 ± 45 Hz, mean difference= -13 Hz, Hedges g = -0.28, CI_95_ = [-0.42, -0.17], N = 20, Figure 4D), again suggesting that the toxin was acting on conductances other than K_V_2.1.

In cells in which K_V_2.1 has a significant conducting role, GxTX-1E normally reduces the threshold for depolarising block of action potentials (Guan *et al*., 2013; Liu & Bean, 2014; Kimm *et al*., 2015). However, we failed to detect significant effects of GxTX-1E on the threshold for AP block in either K_V_2.1^OFF^ or control motoneurons (**K_V_2.1^OFF^**: nACSF = 4.6 ± 2.5 nA, GxTX-1E = 4.7 ± 2.8 nA, mean difference = 0.1 nA, Hedges g = -0.02, CI_95_ = [-0.18, 0.30], N = 12, Figure 4 E-F; **control**: nACSF= 4.1 ± 1.9 nA, GxTX-1E = 4.4 ± 2.0 nA, mean difference = 0.3 nA, Hedges g = 0.15, CI_95_ = [-0.01, 0.35], N = 20).

We also measured AP amplitude, AP half-width, and fAHP amplitude. In keeping with the above data showing that motoneuron firing properties were minimally affected by acute GxTX-1E block of K_V_2 channels, we also saw minimal effects on the AP characteristics assessed (**AP amplitude**- **K_V_2.1^OFF^**: nACSF = 82 ± 4 mV, GxTX-1E = 80 ± 7 mV, mean difference = 2 mV, Hedges g = -0.38, CI_95_ = [- 0.94, 0.26], N = 11, Figure 5A-B; **control**: nACSF = 79 ± 4 mV, GxTX-1E = 78 ± 5 mV, mean difference = -1 mV, Hedges g = -0.24, CI_95_ = [-0.55, 0.12], N = 20; **AP ½ width**- **K_V_2.1^OFF^**: nACSF = 0.57 ± 1.39 ms, GxTX-1E = 0.60 ± 0.18 ms, mean difference = 0.02 ms, Hedges g = 0.14, CI_95_ = [-4e- 03, 0.31], N = 11, Figure 5 A,C; **control**: nACSF = 0.62 ± 1.37 ms, GxTX-1E = 0.64 ± 0.14 ms, mean difference = 0.01 ms, Hedges g = 0.12, CI_95_ = [-0.07, 0.24], N = 20; **fAHP amplitude**- **K_V_2.1^OFF^**: nACSF = 15 ± 5 mV, GxTX-1E = 15 ± 6 mV, mean difference = 0.3 mV, Hedges g = 0.05, CI_95_ = [- 0.16, 0.21], N = 11, Figure 5 D; **control**: nACSF = 15 ± 5 mV, GxTX-1E = 16 ± 5 mV, mean difference = 0.1 mV, Hedges g = 0.02, CI_95_ = [-0.09, 0.17], N = 20).

**Figure 5.**
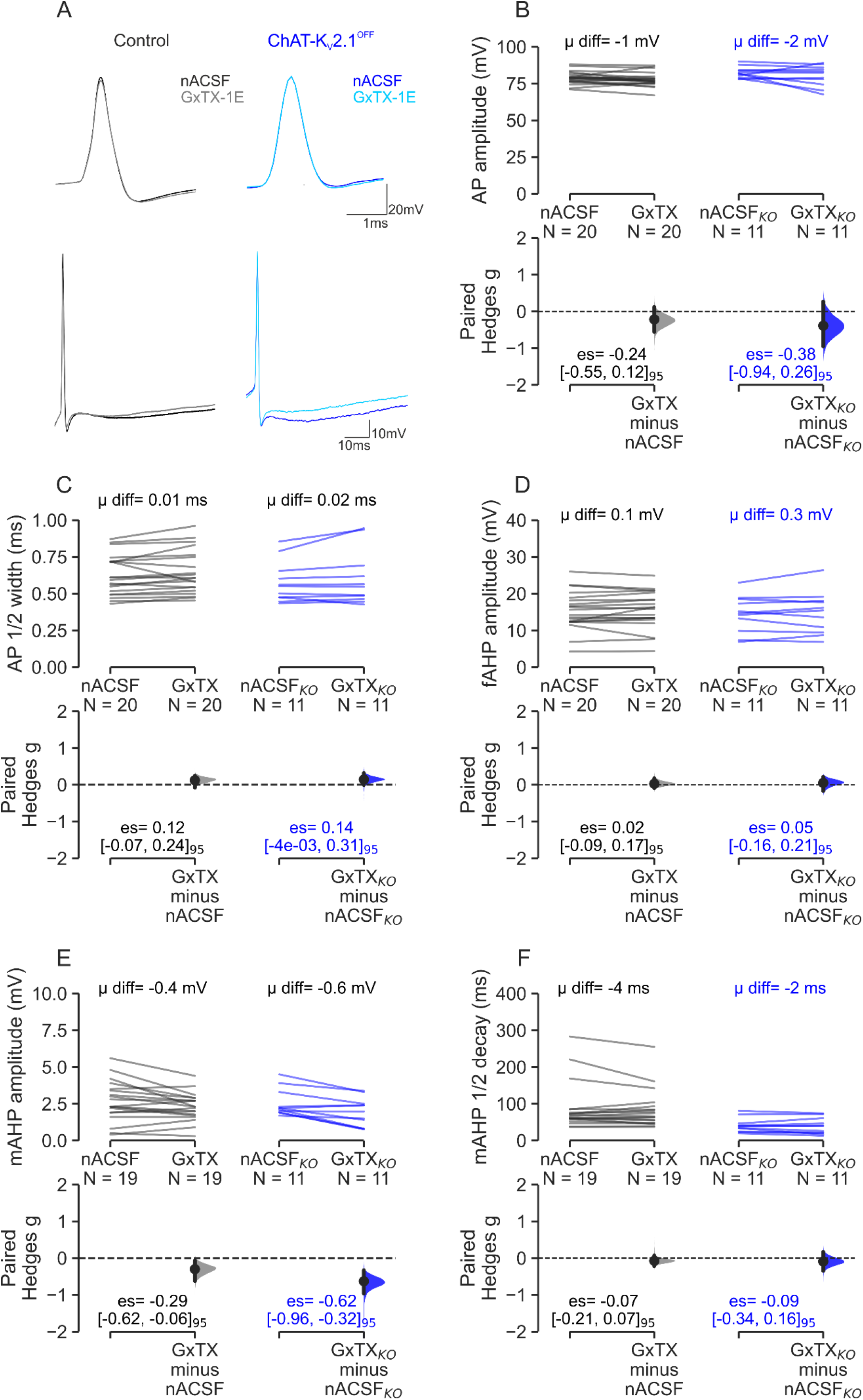
Comparing the effects of K_V_2 inhibition with GxTX-1E on action potential characteristics in control and ChAT-K_V_2.1^OFF^ motoneurons. **(A)** The upper panel shows representative single evoked AP traces from control (left) and ChAT-K_V_2.1**^OFF^** motoneurons (right). The dark colours (black, blue) show spike morphology in the presence of nACSF only and the light colours (grey, sky blue) show morphology after 10 minute perfusion with 100 nM GxTX-1E. The lower panels show longer sweeps in order to visualise the mAHP. **(B-F)** Paired Hedges g for control (left) and ChAT-K_V_2.1^OFF^ (right) motoneurons are shown in Cumming paired estimation plots. Individual motoneurons are plotted on the upper graphs, with each paired set of observations (nACSF followed by 100 nM GxTX-1E) connected by a line. On the lower plots, effect sizes (Hedges g) are shown with bootstrapped sampling distribution (5000 reshuffles). Effect sizes are depicted as dots; 95% confidence intervals are indicated by the vertical error bars. The bootstrapped mean differences (μ diff) are shown on the upper plots, and the effect sizes (es) and 95% confidence intervals [lower, upper] are displayed on the lower plots. **(B)** shows the spike amplitude, **(C)** is action potential ½ width, **(D)** the fAHP amplitude, **(E)** is the mAHP amplitude, and **(F)** is the mAHP ½ decay time. Experimental unit (N) = motoneurons. Number of animals used was as follows: control= 13 (6 females, 7 males), ChAT-K_V_2.1**^OFF^ =** 6 (4 females, 2 males).

Recent evidence suggests K_V_2.1 conductances may contribute to the mAHP (Nascimento *et al*., 2020). In agreement, we found that GxTX-1E had medium and small effects on mAHP amplitudes in K_V_2.1^OFF^ (nACSF = 2.5 ± 0.9 mV, GxTX-1E = 1.9 ± 1.0 mV, mean difference = -0.6 mV, Hedges g = -0.62, CI_95_ = [-0.96, -0.32], N = 11, Figure 5 E) and control motoneurons respectively (nACSF = 2.7 ± 1.4 mV, GxTX-1E = 2.3 ± 1.0 mV, mean difference = -0.4 mV, Hedges g = -0.29, CI_95_ = [-0.62, - 0.06], N= 20). There were no effects on mAHP half decay (**K_V_2.1^OFF^**: nACSF = 41 ± 19 ms, GxTX-1E = 39 ± 22 ms, mean difference = 2 ms, Hedges g = -0.09, CI_95_ = [-0.34, 0.16], N = 11, Figure 5F; **control**: nACSF= 88 ± 65 ms, GxTX-1E = 84 ± 53 ms, mean difference = -4 ms, Hedges g = -0.07, CI_95_ = [-0.21, 0.07], N = 20). Because GxTX-1E-mediated decreases in mAHP amplitude were seen in both control and ChAT-K_V_2.1^OFF^ motoneurons, it is likely that GxTX-1E acted on conductances other than K_V_2.1, albeit with minimal effects.

In summary, apart from small increases in rheobase, and decreases in frequency of the 1^st^ interspike interval and mAHP amplitude in both K_V_2.1^OFF^ and control motoneurons, GxTX-1E had little effect on firing capabilities or AP characteristics. Importantly, GxTX-1E effects were seen in both genotypes, suggesting that the toxin was acting on conductances other than K_V_2.1 (possibly K_V_2.2, see below).

### K_V_2 block increases maximum output and excitability in cortical pyramidal neurons

The results so far suggest that under resting conditions (no C-bouton activation), K_V_2.1 does not play a significant role in regulating motoneuron firing in motor mature (P13-21) mice. Because we were concerned that these results were not consistent with several studies in younger animals (P2-P12) suggesting that K_V_2.1 plays an integral role in regulating motoneuron firing (Fletcher *et al*., 2017; Romer *et al*., 2019; Nascimento *et al*., 2020), we proceeded with a positive control by testing our protocol on layer V cortical pyramidal neurons, in which K_V_2.1 channels have significant electrophysiological functions (Guan *et al*., 2013; Bishop *et al*., 2015; Newkirk *et al*., 2022).

We found that perfusion of cortical slices with GxTX-1E resulted in significant changes to individual pyramidal neuron action potential characteristics with large effect sizes (Figure 6 - figure supplement 1). As demonstrated in similar studies assessing the effect of inhibiting K_V_2.1 in cortical and other brain neurons (Guan *et al*., 2007; Guan *et al*., 2013; Bishop *et al*., 2015; Kimm *et al*., 2015; Newkirk *et al*., 2022), we found large decreases in **spike amplitude** (nACSF = 83 ± 5 mV, N = 7; GxTX-1E = 76 ± 9 mV, N =7, mean difference = -8 mV, Hedges g = -0.97, CI_95_ = [-1.49, -0.45], Figure 6 – figure supplement 1A-B), **AP ½ width** (nACSF = 1.1 ± 0.1 ms, N = 7; GxTX-1E = 1.2 ± 0.1 ms, N = 7, mean difference = 0.1 ms, Hedges g = 1.24, CI_95_ = [0.49, 2.13], Figure 5 - figure supplement 1C), and **fAHP** (nACSF = 6 ± 3 mV, N = 7; GxTX-1E = 4 ± 2 mV, N = 7, mean difference = -3 mV, Hedges g = -0.94, CI_95_ = [-1.77, -0.50], Figure 6 - figure supplement 1D). GxTX-1E also caused a small decrease in **mAHP amplitude** (nACSF = 2.0 ± 0.5 mV, N = 7; GxTX-1E = 1.8 ± 0.6 mV, N = 7, mean difference = -0.20 mV, Hedges g = -0.36, CI_95_ = [-0.96, -0.18], Figure 6 - figure supplement 1E), but **mAHP 1/2 decay** was not affected (nACSF = 86 ± 23 ms, N = 7; GxTX-1E = 79 ± 14 ms, N = 7, mean difference = -12 ms, Hedges g = -0.38, CI_95_ = [-1.39, 0.45], Figure 6 - figure supplement 1F).

GxTX-1E also had effects on repetitive firing of cortical neurons, with increases in the **overall ƒ-I slope** (nACSF = 41 ± 10 mV, N = 7; GxTX-1E = 54 ± 18 mV, N = 7, mean difference = 13 Hz/nA, Hedges g = 0.84, CI_95_ = [0.44, 1.53], Figure 6A-C) and the **initial interval ƒ-I slope** (nACSF = 99 ± 38 mV, N = 7; GxTX-1E = 157 ± 52 mV, N=7, mean difference = 59 Hz/nA, Hedges g = 1.20, CI_95_ = [0.56, 2.07], Figure 6D). There was no consistent effect on firing frequencies, though: both the **overall** (nACSF = 44 ± 6 Hz, N = 7; GxTX-1E = 46 ± 5 Hz, N = 7, mean difference = 2 Hz, Hedges g = 0.35, CI_95_ = [-0.49, 1.32], Figure 5E) and **initial** (nACSF = 113 ± 20 Hz, N = 7; GxTX-1E = 118 ± 12 Hz, N = 7, mean difference = 5 Hz, Hedges g = 0.26, CI_95_ = [-0.72, 1.16], Figure 6F) frequencies were similar in both conditions. This resulted from the neurons reaching depolarising block at significantly lower current thresholds (nACSF = 2.0 ± 0.6 mV, N = 7; GxTX-1E = 1.3 ± 0.4 mV, N = 7, mean difference = -0.6 nA, Hedges g = -1.11, CI_95_ = [-1.78, -0.72], Figure 6G).

**Figure 6.**
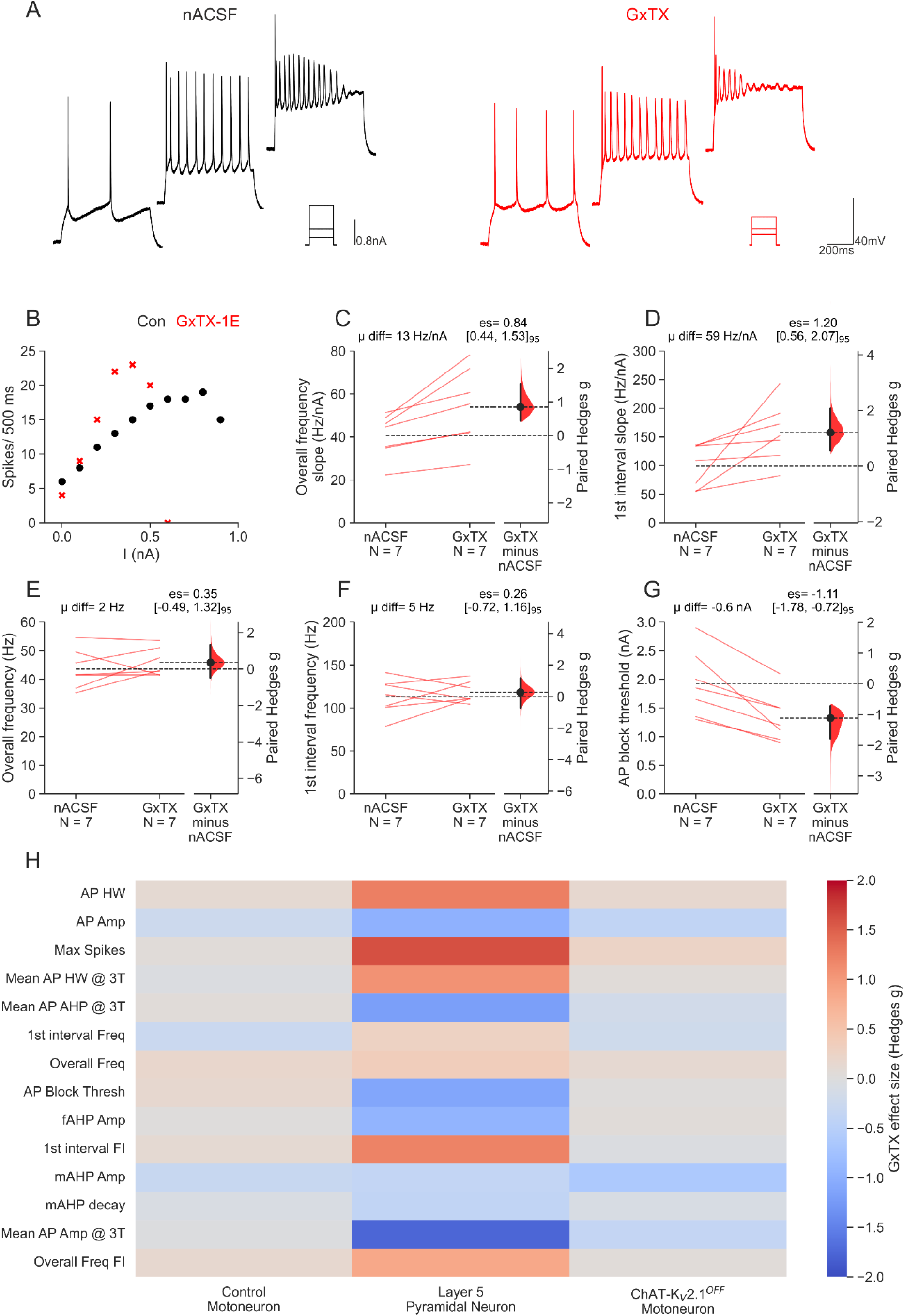
K_V_2 inhibition increases excitability of cortex layer V pyramidal neurons. **(A)** Representative traces showing pyramidal neuron firing in response to 500 ms current steps of increasing magnitude before (nACSF, black) and after (GxTX-1E, red) GxTX-1E perfusion. **(B)** Scatter plot of number of spikes in response to increasing current input in a representative pyramidal neuron in nACSF (black circles) and after GxTX-1E perfusion (red crosses). **(C-G)** Cumming paired estimation plots for individual neurons plotted on the left axes with each paired set of observations (nACSF followed by 100 nM GxTX-1E) connected by a line. On the right axes, effect sizes (Hedges g) are plotted as a bootstrapped sampling distribution (5000 reshuffles). Effect sizes are depicted as dots; 95% confidence intervals are indicated by the vertical error bars. The bootstrapped mean differences (μ diff) are shown on the upper plots, and the effect sizes (es) and 95% confidence intervals [lower, upper] are displayed on the lower plots. Experimental unit (N) = neurons recorded from 3 animals (2 males, 1 female). Plots show overall frequency slope **(C)**, 1^st^ interval frequency slope **(D)**, maximum overall frequency **(E)**, maximum 1^st^ interval frequency **(F)**, and AP block threshold **(G). (H)** Heat map showing the effect sizes (Hedges g) for GxTX-1E induced changes in action potential and firing characteristics in mature control motoneurons, layer 5 cortical pyramidal neurons, and mature ChAT-K_V_2.1^OFF^ motoneurons. Dark colours represent large positive or negative effect sizes.

In summary, we found that GxTX-1E block of K_V_2 channels had little effect on the firing capabilities of mature control and K_V_2.1^OFF^ motoneurons. But the same experiments in layer 5 cortical pyramidal neurons greatly increased excitability and maximum firing, in line with previous studies. The clear difference in paired sample effect sizes (Hedges g) on the parameters assessed suggests that, compared to pyramidal neurons, K_V_2 channels (both 2.1 and 2.2) play only a minor role in regulating motoneuron repetitive firing (Figure 5H).

### Postnatal clustering does not significantly alter _KV_2 regulation of motoneuron firing

During post-natal motoneuron development, K_V_2.1 channels organise into high density macroclusters (Wilson *et al*., 2004). It has been proposed that, like in other neurons (Misonou *et al*., 2004; Misonou *et al*., 2005), channel clustering modulates K_V_2.1 conductances in motoneurons (Romer *et al*., 2014; Romer *et al*., 2019). Because our patch clamp experiments were done at an age (post-natal week 3) at which K_V_2.1 clusters appear mature (Wilson *et al*., 2004), we investigated whether development of channels might contribute to differences in our results compared to previous studies (Fletcher *et al*., 2017; Romer *et al*., 2019; Nascimento *et al*., 2020). To do this, we first used immunohistochemistry to quantitatively assess the expression and membrane localisation of K_V_2.1 channels in lumbar motoneurons from mice aged P2-P3 (neonatal, Figure 7A-A3), P6-P7 (transition) & P21 (motor mature, Figure 7B-B3). There was a significant increase in the density of VAChT^ON^ C-boutons with development (R_s_ = 0.49, p = 1e-09, N =137, Spearman’s rank, Figure 7C), while K_V_2.1 density (# puncta pre 100µm^2^) showed a weak negative relationship with age (R_s_ = -0.17, p = 0.042, N = 137, Figure 7D). As qualitative data previously suggested (Wilson *et al*., 2004), we found a significant positive relationship between the percentage of K_V_2.1 localised opposite C-boutons and age (R_s_ = 0.73, p = 2e-23, N = 137, Figure 7E-F1). These data show that on neonatal motoneurons, K_V_2.1 channels are mainly organised in a dispersed microcluster configuration (Figure 7F), and many are not associated with C-boutons (Figure 7F1). That is, as postnatal development progresses and motoneurons (Smith & Brownstone, 2020) and motor behaviour mature (Altman & Sudarshan, 1975), large macroclusters of K_V_2.1 channels form opposite C-bouton synapses.

**Figure 7.**
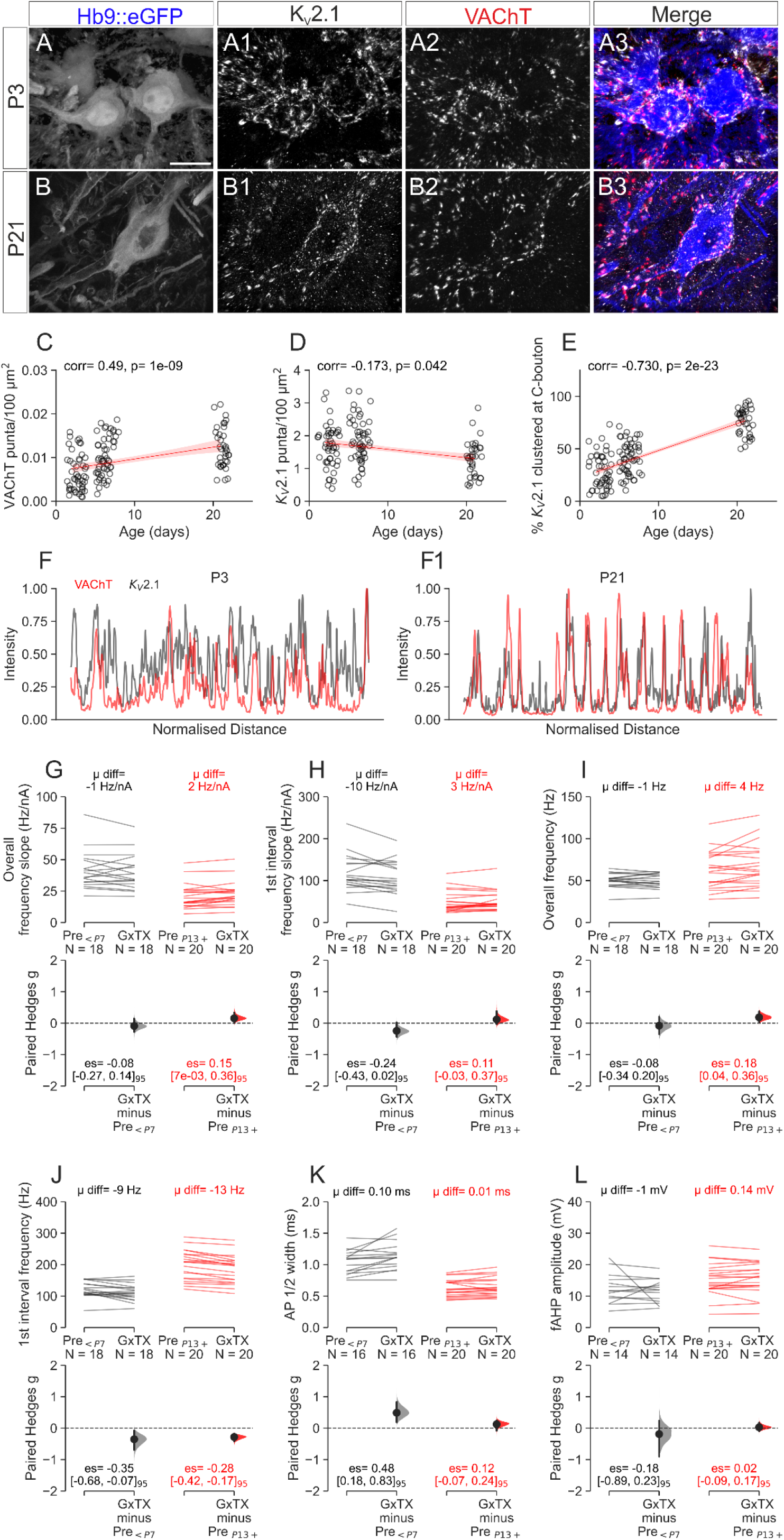
Developmental clustering of K_V_2.1 channels does not reduce their contribution to motoneuron firing. 3D projections of Z-stacks taken at 60x on a confocal microscope showing raw GFP expression under the Hb9 promotor **(A, B)**, immuno-labelling of K_V_2.1 (**A1-B1)** and VAChT (**A2-B2**, depicting presynaptic C-boutons). Images merged in **(A3-B3)** using blue, white, and red, respectively. **(C-E)** Scatter plots with regression models fitted (red lines, with shaded area showing bootstrapped 95% confidence intervals, 5000 resamples) for the density of VAChT^+^ C-boutons **(C)**, density of K_V_2.1 **(D)**, and % K_V_2.1clustered to C-boutons on individual motoneurons from throughout development **(E)**. Note that measurements were taken at P2-3, P6-7 and P21, however jitter was added to the x axes to avoid excessive overlap. A total of 137 motoneurons were sampled from 8 animals (**P2-3**: 3 mice, N=48; **P6-7**: 3 mice N=51; **P21**: 2 mice, N=36; all females). Spearman’s rank was used to determine correlation coefficients and p values. **(F-F1)** Intensity plots (intensity/maximum intensity) showing the distribution of VAChT and K_V_2.1 signal around the perimeter of a 60x confocal image (1μm section) of a representative motoneuron from a P3 (**F**) and P21 (**F1**) mouse. **(G-L)** Paired Hedges g for overall frequency slope **(G)**, 1^st^ interval frequency slope **(H)**, maximum overall frequency **(I)**, maximum 1^st^ interval frequency **(J)**, AP ½ width **(K)**, and fAHP **(L)** in young (P2-7, left) and motor mature (P13-21, right) motoneurons are shown in Cumming paired estimation plots. Individual motoneurons are plotted on the upper graphs, with each paired set of observations (nACSF followed by 100 nM GxTX-1E) connected by a line. On the lower plots, effect sizes (Hedges g) are plotted with bootstrapped sampling distribution (5000 reshuffles). Effect sizes are depicted as dots; 95% confidence intervals are indicated by the vertical error bars. The bootstrapped mean differences (μ diff) are shown on the upper plots, and the effect sizes (es) and 95% confidence intervals [lower, upper] are displayed on the lower plots. Experimental unit (N) shown on each plot. The number of motoneurons and animals used for each age was as follows: P2-3 = 8 cells from 4 mice (2 males, 2 females); P6-7 = 10 cells from 5 mice (2 males, 3 females; 9 animals total for the <P7 group), for mature motoneurons (P13+) 13 were mice were used (6 females, 7 males). Scale bar in **A** = 20μm.

To determine whether development of K_V_2.1 channels alters motoneuron electrical properties, we compared the effects of GxTX-1E on immature and mature motoneuron firing and AP characteristics. As with mature motoneurons (described above), GxTX-1E had minimal effect on **overall ƒ-I slope** (nACSF = 41 ± 15 Hz/nA, N = 18; GxTX-1E = 40 ± 14 Hz/nA, N = 18, mean difference = -1 Hz/nA, Hedges g = -0.08, CI_95_ = [-0.27, -0.14], Figure 7 G), **1^st^ interval ƒ-I slope** (nACSF = 118 ± 43 Hz/nA, N = 18; GxTX-1E = 107 ± 39 Hz/nA, N = 18, mean difference = 10 Hz/nA, Hedges g = -0.24, CI_95_ = [-0.43, 0.02], Figure 7 H), **overall frequency** (nACSF = 50 ± 8 Hz, N = 18; GxTX-1E = 49 ± 9 Hz, N = 18, mean difference = -1 Hz, Hedges g = -0.08, CI_95_ = [-0.34, 0.20], Figure 7 I) or **1^st^ interval frequency** (nACSF = 120 ± 25 Hz, N = 18; GxTX-1E = 111 ± 26 Hz, N = 18, mean difference = -9 Hz, Hedges g = -0.35, CI_95_ = [-0.68, -0.07], Figure 7 J). Despite the lack of effects on firing characteristics, we did find that GxTX-1E increased the AP ½ width by 0.10 ms in young compared to 0.01 ms in mature motoneurons (young: nACSF = 1.0 ± 0.2 ms, N = 16; GxTX-1E = 1.1 ± 0.2 ms, N = 16, mean difference = 0.10 ms, Hedges g = 0.48, CI_95_ = [0.18, 0.83], Figure 7K). As with mature motoneurons, there was no effect of GxTX-1E on fAHP amplitude in young motoneurons (nACSF = 13 ± 5 mV, N = 14; GxTX-1E = 12 ± 4 mV, N = 14, mean difference = -1 mV, Hedges g = -0.18, CI_95_ = [-0.89, 0.23], Figure 7 L). Together, these results show that K_V_2 conductances have a significant effect on AP repolarisation in neonatal motoneurons.

Overall, the developmental anatomy and physiology data suggest that the lack of effect on motoneuron firing of both the chronic knockout of K_V_2.1 channels and acute block of K_V_2 channels by GxTX-1E is not due to developmental differences or channel clustering.

### Muscarinic increase in excitability is preserved in K_V_2.1^OFF^ motoneurons

Motoneuron K_V_2.1 channels are opposed to, and thought to be modulated by C-boutons via M2 muscarinic acetylcholine receptors (Muennich & Fyffe, 2004; Nascimento *et al*., 2020). Our data to this point suggest that the conducting role for K_V_2.1 is minimal in the absence of muscarinic activation. To assess whether K_V_2.1 conductances play a role in mediating the effects of C-boutons, we proceeded to compare motoneuron responses to muscarine in control and ChAT-K_V_2.1^OFF^ mice (Figure 8).

**Figure 8.**
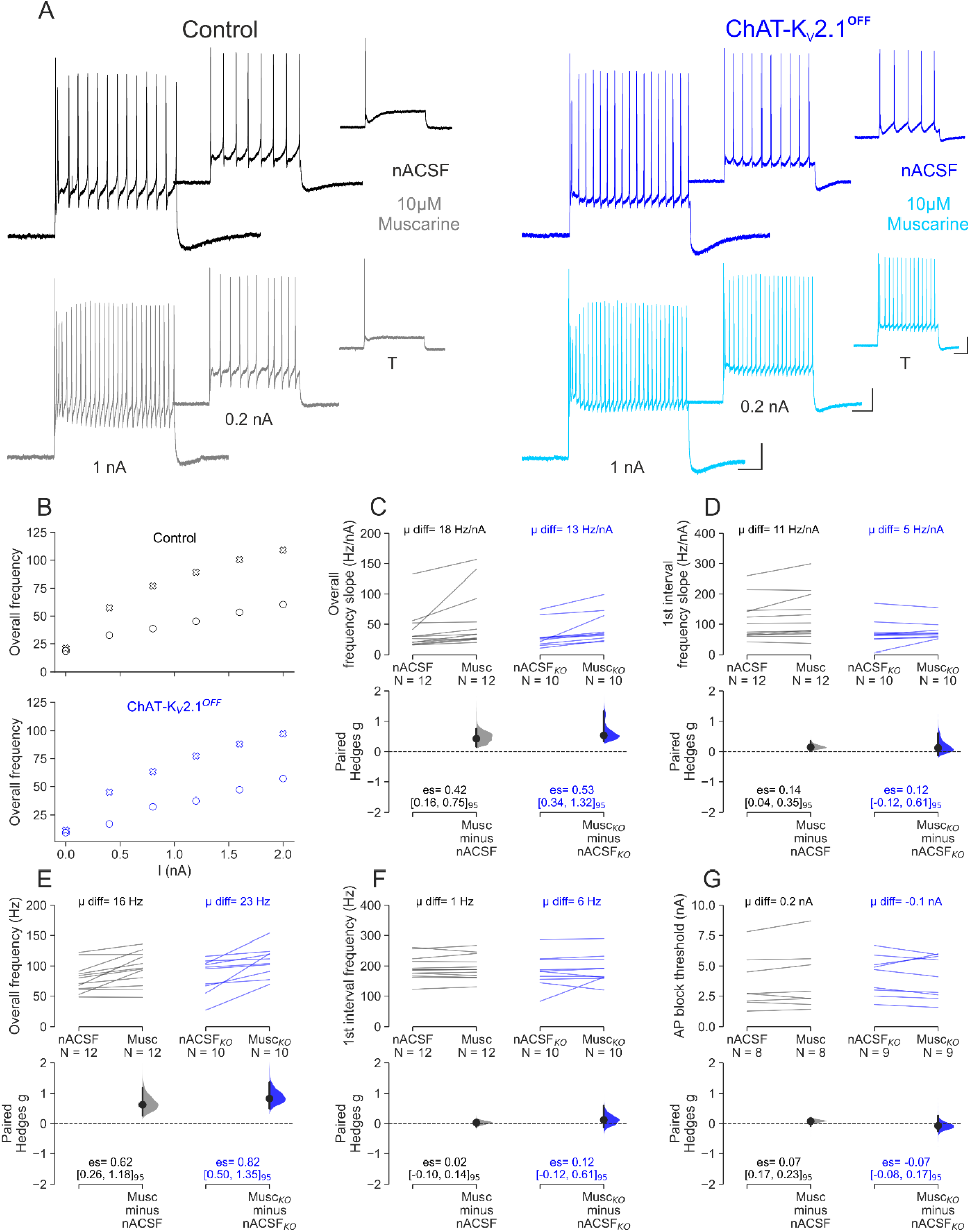
Muscarine increases motoneuron excitability despite K_V_2.1 cKO. **(A)** The changes in firing characteristics following perfusion with 10µM muscarine were compared for motoneurons of control (left) and ChAT- K_V_2.1 ^OFF^ mice (right). Representative traces at threshold (T), 0.2 nA and 1nA above threshold before (nACSF, black and dark blue traces) and after perfusion with muscarine (10 μM, grey and sky blue). **(B)** Scatter plots showing the overall frequency in response to increasing current inputs in control (upper) and ChAT- K_V_2.1 ^OFF^ (lower) motoneurons. **(C-H)** Paired Hedges g for overall frequency slope **(C)**, 1^st^ interval frequency slope **(D)**, maximum overall frequency **(E)**, maximum 1^st^ interval frequency **(F)**, and AP block threshold **(G)** in control (left) and ChAT-K_V_2.1^OFF^ (right) motoneurons are shown in Cumming paired estimation plots. Control (left) and ChAT- K_V_2.1^OFF^ (right) motoneurons are shown in Cumming paired estimation plots. Individual motoneurons are plotted on the upper graphs; each paired set of observations (nACSF followed by 10 μM Muscarine) connected by a line. On the lower plots, effect sizes (Hedges g) are plotted as a bootstrapped sampling distribution (5000 reshuffles). Effect sizes are depicted as dots; 95% confidence intervals are indicated by the vertical error bars. The bootstrapped mean differences (μ diff) are shown on the upper plots, and the effect sizes (es) and 95% confidence intervals [lower, upper] are displayed on the lower plots. Experimental unit (N) = motoneurons. Number of animals used was as follows: control= 8 (2 females, 6 males), ChAT-K_V_2.1**^OFF^** = 6 (2 females, 4 males). Scale bars in **A-B**: horizontal= 100 ms, vertical= 20 mV.

In neonatal motoneurons, exogenously applied muscarine has been reported to have mixed effects due to the different sub-types (M2/M3) of receptors (Nascimento *et al*., 2019). In agreement with those findings, we found that P2-P7 motoneurons (n=9) had large depolarisations (mean diff = 9.5 mV) in response to muscarine (10 uM), whereas in motoneurons from P13 and older mice (n=10), muscarine did not affect resting membrane potential (mean diff = -0.5 mV, Figure 8- figure supplement 1). Rather, the effects of muscarine were dominated by those shown to be due to M2 receptor activation (Miles *et al*., 2007) as outlined below.

Perfusion of control and K_V_2.1^OFF^ motoneurons with 10 µM muscarine had little effect on **AP amplitude** (**Control:** mean difference = -4 mV, Hedges g = -0.57, CI_95_ = [-1.04, 0.12], N =11; **K_V_2.1^OFF^**: mean difference = -0.1 mV, Hedges g = -5e-03, CI_95_ = -0.19, 0.25], N = 9, Figure 8 - figure supplement 2 A-B), **½ width** (**Control:** mean difference = 6e-03 ms, Hedges g = 0.04, CI_95_ = [-20, 0.33], N =11; **K_V_2.1^OFF^**: mean difference = -0.03 ms, Hedges g = -0.26, CI_95_ = [-0.65, -0.02], N = 9, Figure 8 - figure supplement 2 C), or **fAHP amplitude** (**Control:** mean difference = -0.2 mV, Hedges g = -0.06, CI_95_ = [-0.48, 0.21], N =11; **K_V_2.1^OFF^**: mean difference = 0.2 mV, Hedges g = 0.02, CI_95_ = [-0.14, 0.07], N = 9, Figure 8 - figure supplement 2 D). However, the **mAHP amplitude** (**Control:** mean difference = -1.7 mV, Hedges g = -1.14, CI_95_ = [-1.7, -0.63], N =11; **K_V_2.1^OFF^**: mean difference = -3.0 mV, Hedges g = -2.5, CI_95_ = [-3.82, -1.53], N =8, Figure 8 - figure supplement 2 E) and **½ decay time Control:** mean difference = -25 ms, Hedges g = -1.26, CI_95_ = [-2.08, -0.56], N =11; **K_V_2.1^OFF^**: mean difference = -51 ms, Hedges g = -2.1, CI_95_ = [-3.36, -1.32], N =8, Figure 8 - figure supplement 2 F) were significantly decreased in both control and K_V_2.1^OFF^ motoneurons.

In agreement with previous work in control motoneurons (Miles *et al*., 2007), and in line with the reduction in mAHP conductances, muscarine increased **overall ƒ-I slope** (nACSF = 38 ± 33 Hz/nA, Muscarine = 56 ± 48 Hz/nA, mean difference = 18 Hz/nA, Hedges g = 0.42, CI_95_ = [0.16, 0.75], N =12, Figure 8C), but not the **ƒ-I slope of the first interval** (nACSF = 113 ± 68 Hz/nA, Muscarine = 124 ± 78 Hz/nA, mean difference = 11 Hz/nA, Hedges g= 0.14, CI_95_=[0.04, 0.35], N = 12, Figure 8D). In addition, muscarine induced moderate increases in **overall firing frequency** (nACSF = 79 ± 24 Hz, Muscarine = 95 ± 27 Hz, mean difference = 16 Hz, Hedges g = 0.62, CI_95_ = [0.26, 1.18], N = 12, Figure 8E). The 1^st^ spike interval frequency was not altered by muscarine in control motoneurons (nACSF = 194 ± 40 Hz, Muscarine = 195 ± 40 Hz, mean difference = 1 Hz, Hedges g = 0.02, CI_95_ = [0.10, 0.14], N = 12, Figure 8F).

Muscarine was similarly effective on K_V_2.1^OFF^ motoneurons. The **overall ƒ-I slope** was increased (nACSF = 11 ± 22 Hz/nA, Muscarine = 44 ± 26 Hz/nA, mean difference = 13 Hz/nA, Hedges g = 0.53, CI_95_ = [0.34, 1.32], N = 10, Figure 8C), while the **ƒ-I slope of the first interval** was unchanged (nACSF = 72 ± 43 Hz/nA, Muscarine = 77 ± 30 Hz/nA, mean difference = 5 Hz/nA, Hedges g = 0.12, CI_95_ = [-0.12, 0.61], N = 10, Figure 8D). There was a large increase in the **overall frequency** (nACSF = 84 ± 28 Hz, Muscarine = 107 ± 24 Hz, mean difference = 23 Hz, Hedges g = 0.82, CI_95_ = [0.50, 1.35], N = 10, Figure 8E). As with the control motoneurons, **initial frequency** was not altered by muscarine in ChAT-K_V_2.1^OFF^ motoneurons (nACSF = 183 ± 54 Hz, Muscarine = 189 ± 47 Hz, mean difference = 6 Hz, Hedges g = 0.12, CI_95_ = [-0.12, 0.61], N = 10, Figure 8F).

As K_V_2 channels have been shown in other neurons to prevent depolarising block to maintain firing during high synaptic drive, we asked whether muscarinic receptor activation might modulate K_V_2.1 to increase the current threshold at which spike output is blocked. However, muscarine had no effect on depolarising block threshold in either control (nACSF = 3.6 ± 2.2 nA, Muscarine = 3.8 ± 2.5 nA, mean difference = 0.2 nA, Hedges g = 0.07, CI_95_ = [0.17, 0.23], N = 12) or K_V_2.1^OFF^ motoneurons (nACSF = 4.2 ± 1.7 nA, Muscarine = 4.1 ± 1.8 nA, mean difference = -0.1 nA, Hedges g = 0.07, CI_95_ = [-0.08, 0.17], N = 10, Figure 8G).

Taken together, these results show that K_V_2.1 is not required for muscarine-evoked increases in motoneuron excitability.

### Motor behaviour and C-bouton amplification is preserved in ChAT-K_V_2.1^OFF^mice

Despite our in vitro data suggesting that K_V_2.1 channels play minimal conducting role in motor amplification, we reasoned that cKO mice would still show deficits in high force tasks if they had a significant non-conducting role. To determine whether motor function was impaired in cKO mice, we performed several behavioural assays that rely on high force production. Voluntary motor behaviour over a chronic period was assessed by individually housing male (Figure 9 - figure supplement 1A) and female (Figure 9 - figure supplement 1B) mice of both genotypes in cages with *ad libitum* access to a running wheel for 23 days. Male control (N = 5) and ChAT-K_V_2.1^OFF^ (N = 6) mice ran similar daily distances (two-way repeated measures ANOVA, genotype, F = 0.11, p = 0.74; day, F = 0.68, p = 0.85; interaction, F = 0.14, p = 0.99) over the 23 day period. For females, ChAT-K_V_2.1^OFF^ mice seemed to lack a training response, whereas control females showed an increase in distance over the 23 days (genotype, F = 5.6, p = 0.01; day, F = 1.5, p = 0.043; interaction, F = 1.27, p = 0.15).

We next examined the running capacity of mice on a treadmill inclined 15° to increase locomotor force demands. The maximum speed for male ChAT-K_V_2.1^OFF^ group was on average 6 cm/s slower than controls (control = 46 ± 6 cm/s, N = 7; ChAT-K_V_2.1^OFF^ = 41 ± 6 cm/s, N = 9, mean difference = - 6 cm/s, Hedges g = -0.92). Although the effect size was large, the 95% confidence interval was wide and crossed 0 (CI_95_ = [-1.84, 0.07]), indicating a relatively low degree of certainty in the magnitude of this effect (Figure 9 - figure supplement 1C). The maximum distance run at 60% of maximum speed (control = 0.23 ± 0.07 km, N = 8; ChAT-K_V_2.1^OFF^ = 0.22 ± 0.10 km, mean difference = -0.01 km, N = 9, Hedges g = -0.12, CI_95_ = [-1.14, 0.91], Figure 9 - figure supplement 1D) and grip strength (control = 10 ± 1.4 g/g body weight, N = 5; ChAT-K_V_2.1^OFF^ = 10 ± 1.4 g/g body weight, mean difference = 0.5 g/g body weight, N = 7, Hedges g = 0.35, CI_95_ = [-1.42, 1.42], Figure 9 - figure supplement 1E) were similar in both genotypes for male mice.

In female mice, there were no differences between the groups for maximum speed (control = 51 ± 6 cm/s, N = 8; ChAT-K_V_2.1^OFF^ = 48 ± 5 cm/s, mean difference = -3 cm/s, Hedges g = -0.53, CI_95_ = [- 1.73, 0.53], N = 8) or distance run on a treadmill at 60% maximum speed (control = 0.39 ± 0.18 km, N = 8; ChAT-K_V_2.1^OFF^ = 0.50 ± 1.97 km, N = 8, mean difference = 0.11 km, Hedges g = 0.53, CI_95_ = [-0.50, 1.70], Figure 9 - figure supplement 1G). Grip strengths were also similar (control = 12 ± 1 g/g BW, N = 6; ChAT-K_V_2.1^OFF^ = 12 ± 2 g/g BW, N = 12, mean difference = 0.48 g/g BW, Hedges g = - 0.13, CI_95_ = [-1.26, 0.84], Figure 9 - figure supplement 1H). Thus, we were unable to detect any significant deficits in either male or female ChAT-K_V_2.1^OFF^ mice using these particular tasks.

C-bouton mediated amplification can be more specifically assessed *in vivo* by comparing EMG amplitudes during walking and swimming: when C-bouton synaptic transmission is genetically silenced, the ratio of the EMG amplitude during swimming vs walking is reduced compared to mice with functional C-bouton signalling (Zagoraiou *et al*., 2009; Landoni *et al*., 2019; Konsolaki *et al*., 2020). Thus, if K_V_2.1 were critical to C-bouton function, a similar decrease in motor amplification would be expected following cKO in motoneurons. However, we found no difference in EMG amplification during swimming between control and ChAT-K_V_2.1^OFF^ mice in either the extensor medial gastrocnemius (MG) muscle (control = 4.1 ± 3.6, N = 5, ChAT-K_V_2.1^OFF^ = 6.7 ± 5.5, N = 7, mean difference = 1.25, Hedges g = 0.25, CI_95_ = [-1.11, 1.29], Figure 9A-A1, C) or the flexor tibialis anterior (TA) muscle (control= 2.5 ± 0.9, N=5, ChAT-K_V_2.1^OFF^ = 2.5 ± 0.8, N=7, mean difference= - 0.17, Hedges g= -0.21, CI_95_= [-1.86, 1.25], Figure 9B-B1, C1).

**Figure 9.**
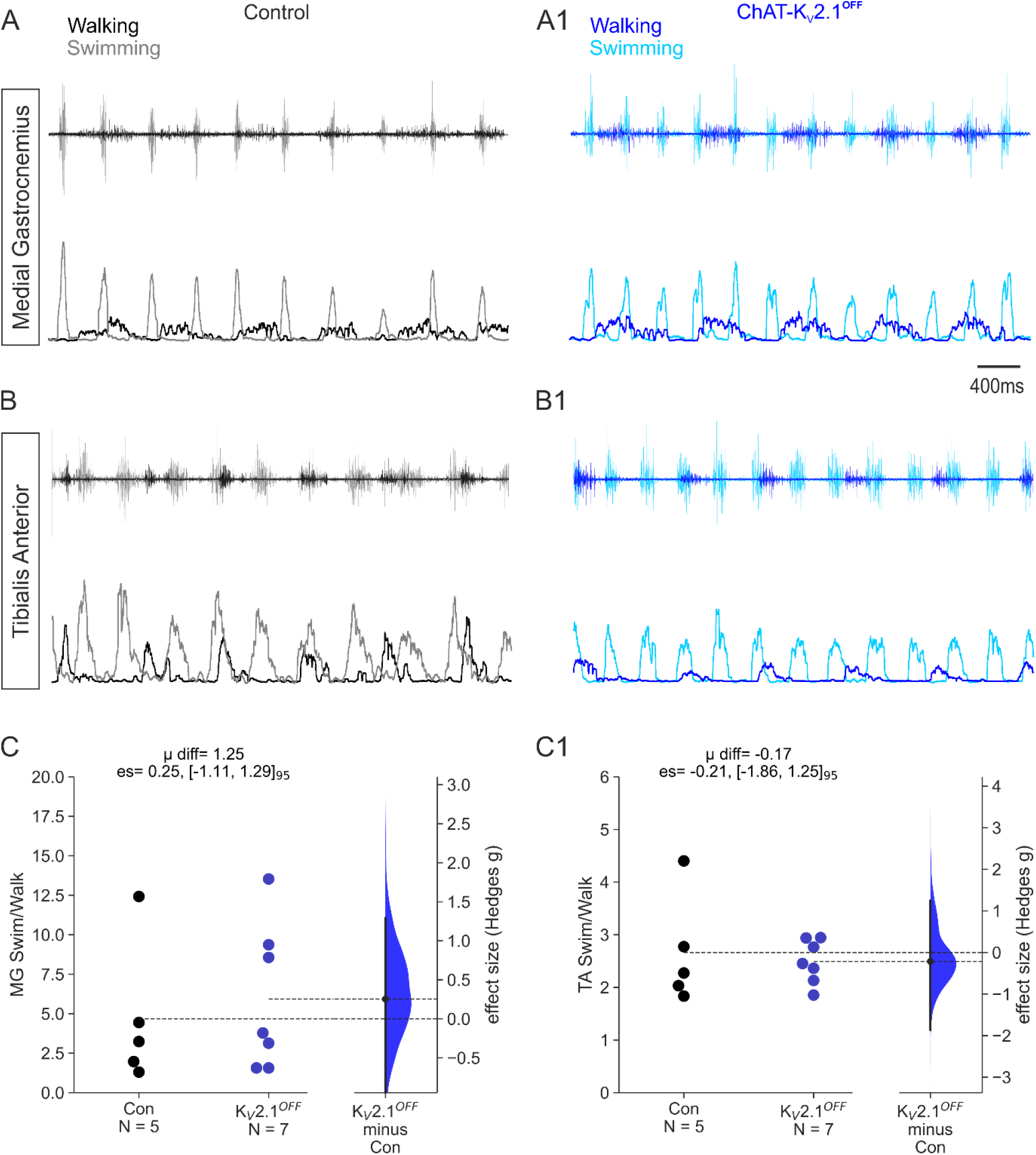
Motoneuron amplification is preserved in ChAT-K_V_2.1^OFF^ mice. Representative medial gastrocnemius (MG) and tibalis anterior (TA) EMG traces from a control **(A-B)** and ChAT-K_V_2.1^OFF^ mouse **(A1-B1)**. For control mice **(A-B)**, black traces are those recorded during walking at 0.15 m/s and grey traces are during swimming. In ChAT-K_V_2.1^OFF^ mice, blue traces are walking at 0.15 m/s and sky blue is swimming. For each muscle, top traces are raw EMGs and lower traces are corresponding RMS amplitudes. **(C-C1)** Gardner-Altman estimation plots for MG and TA. Control and ChAT-K_V_2.1**^OFF^** groups are plotted the left axes and the bootstrapped sampling distribution (5000 reshuffles) for Hedges g effect sizes are plotted on the right. Each point represents mean ratio for both MG or TA muscles. The Hedges g effect size is depicted as a dot; the 95% confidence interval is indicated by the vertical error bar. The mean difference (μ diff), effect sizes (es) and 95% confidence intervals [lower, upper] are displayed at the top of each plot. Experimental unit (N) = animals, all were females.

In summary, the only behavioural effect of eliminating K_V_2.1 from cholinergic neurons that we observed was that of a possible training effect in female mice given *ad libitum* access to a running wheel. Notably, these data suggest that either K_V_2.1 channels are not necessary for motoneuron amplification – the key reported role of C-boutons – or that their absence can be compensated to maintain behavioural homeostasis.

### K_V_2.2 is co-expressed with K_V_2.1 opposite C-bouton synapses

The lack of behavioural deficits and in particular the lack of motoneuron amplification following cKO of K_V_2.1 were surprising findings considering that these channels are prominent at all C-bouton synapses and thought to be an integral part of the synaptic machinery (Muennich & Fyffe, 2004; Romer *et al*., 2019; Nascimento *et al*., 2020). Although K_V_2.1 is considered to be the predominant K_V_2 subunit in motoneurons (Deardorff *et al*., 2021), in many brain neurons K_V_2.1 subunits share some functional homology and are co-expressed with K_V_2.2 subunits (Guan *et al*., 2007; Guan *et al*., 2013; Bishop *et al*., 2015; Kimm *et al*., 2015; Johnson *et al*., 2018; Kirmiz *et al*., 2018). We therefore proceeded with immunohistochemistry experiments to identify whether K_V_2.2 is also expressed in motoneurons. We found that K_V_2.2 puncta were abundant in the spinal cord and often, but not always co-localised with K_V_2.1 (Figure 10A-F). In the dorsal horn (Figure 10 - figure supplement 1 A-C1), K_V_2.1 was predominantly expressed in the deeper laminae, whereas K_V_2.2 positive cells were concentrated in the superficial laminae. In the deep dorsal/intermediate laminae (Figure 10 - figure supplement 1 D-F1), K_V_2.2 expression was mostly confined to medial regions, and as well as forming large clusters on somata, an area of smaller, more diffuse puncta were located lateral to the dorsal columns (Figure 10 - figure supplement 1 D-F1). In the same region, K_V_2.1 puncta were distributed across the medio-lateral axis, mainly in neurons without K_V_2.2 labelling. In the ventral horn (Figure 10A-F), K_V_2.2 seemed to be exclusively confined to motoneurons, where labelling was co-localised with K_V_2.1. Co-labelling experiments with ChAT confirmed that K_V_2.2 clusters are localised to the postsynaptic membrane opposite C-boutons (Figure 10D-F). Thus, K_V_2.2 channels are clearly co- localised with K_V_2.1 channels at motoneuron membranes apposing pre-synaptic C-boutons.

**Figure 10.**
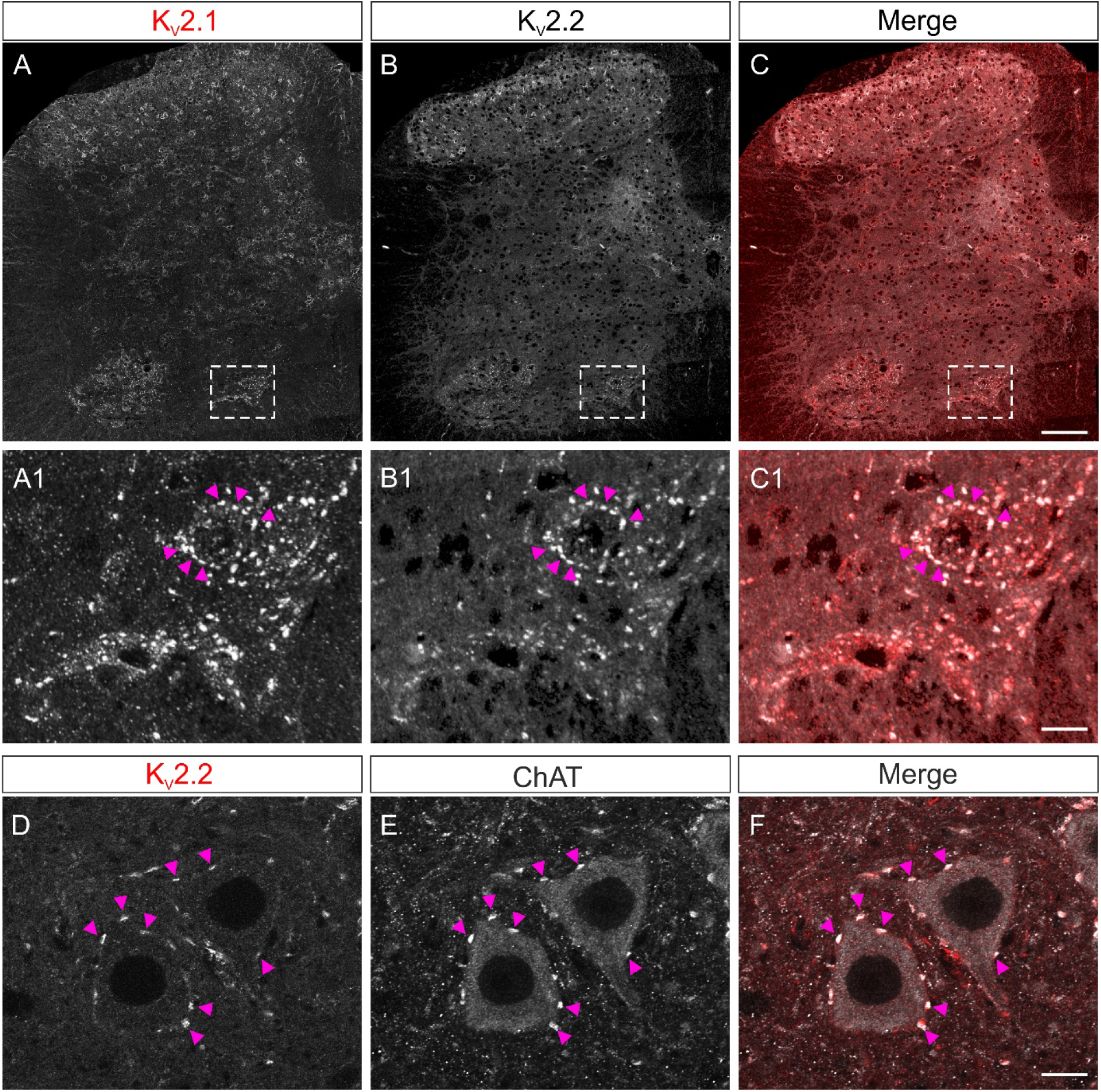
K_V_2.2 is expressed in the spinal cord and co-localises with K_V_2.1 opposite to C-boutons. **(A-C)** 40x confocal tiled, Z stack projection images (5 x 1µm slices) of the superficial laminae of the lumbar spinal cord stained for K_V_2.1 **(A)** and K_V_2.2 **(B)**; merged image shown in **(C)**. **(A1-C1)** Boxes depicted in A-C, cropped and expanded. (**D-F)** 40x confocal single optical sections showing K_V_2.2 **(D)** expression opposite VAChT^+^ C boutons **(E),** with a merge of both channels in **(F)**. Magenta arrows show clustering of K_V_2.2 channels to the C-bouton synapses. Scale bars in **(A-C)** = 80 µm, **(A1-C1)** = 25 µm, **(D-F)** = 10 µm.

## Discussion

K_V_2.1 channels are expressed by spinal motoneurons where they largely aggregate opposite C-bouton synapses (Muennich & Fyffe, 2004), clustering in the post-natal period as the motor system matures (Altman & Sudarshan, 1975; Wilson *et al*., 2004; Smith *et al*., 2017; Smith & Brownstone, 2020). K_V_2.1 delayed rectifier currents have been implicated as crucial regulators of motoneuron firing and C-bouton amplification of motor output (Fletcher *et al*., 2017; Romer *et al*., 2019; Nascimento *et al*., 2020). Here, we sought to define their function using a genetic strategy to eliminate these membrane proteins from cholinergic neurons, including motoneurons. We were somewhat surprised that in doing so, motoneuron physiology and animal behaviour were relatively unaffected. The data presented here thus challenge some current concepts of motoneuron physiology and raise new questions about the role of these prominent channels in motoneuron behaviour. We suggest that the primary function of K_V_2 proteins – K_V_2.1 and K_V_2.2 together – is non-conducting, and that K_V_2.2 is sufficient for this function if K_V_2.1 channels are absent.

### Co-expression of K_V_2.1 and K_V_2.2 in motoneurons

The structure and function of K_V_2 channels has been studied for over 3 decades, since K_V_2.1 was shown to be expressed in rat brain (Frech *et al*., 1989). K_V_2.2 channels were discovered soon after K_V_2.1, and although the two subtypes have similar electrophysiological properties, substantial differences in expression profiles across the brain were initially seen between the two (Hwang *et al*., 1992). Evidence was then presented that not only were these channels differentially expressed in different neurons, but membrane localisation of the two channels was distinct, with some neurons having more evident expression of K_V_2.2 (“CDRK” at that time) in their processes than on their soma (Hwang *et al*., 1993a). Subsequently, K_V_2.1 became the primary focus of further studies of K_V_2 regulation of neuronal excitability.

It later became clear, however, that the previous characterisation of K_V_2.2 was based on an unfortunate cloning error and that both K_V_2.1 isoforms are “co-localised in the somata and proximal dendrites of cortical pyramidal neurons, and are capable of forming heteromeric channels” (Kihira *et al*., 2010). Nonetheless, much of the focus has remained on K_V_2.1 and only recently has K_V_2.2 received increasing attention (Kihira *et al*., 2010; Guan *et al*., 2013; Bishop *et al*., 2015; Johnson *et al*., 2018; Kirmiz *et al*., 2018). But K_V_2.2 has remained unexplored in motoneurons. Here, we ultimately – after seeing minimal effects following elimination of K_V_2.1 – show that K_V_2.2 is also expressed post-synaptically to C-boutons, is co-clustered with K_V_2.1, and as such, should be taken into consideration when investigating C-bouton function. Whether K_V_2.2 expression increases in ChAT-K_V_2.1^OFF^ motoneurons as a compensatory mechanism remains unknown.

### Reconciling differences with published studies assessing K_V_2.1 function in motoneurons

Our experiments were initially performed with the assumptions that K_V_2.1 was the predominant K_V_2 subtype in motoneurons, and that it had a significant conducting role. We expected that K_V_2.1 cKO would reduce delayed rectification and thus alter motoneuron firing characteristics. However, the only difference in mature ChAT-K_V_2.1^OFF^ motoneurons compared to controls was a capacity to fire at higher rates. Interestingly, given the filtering properties of muscle fibres, these high frequencies may not be relevant to muscle contraction (Enoka & Farina, 2021).

If K_V_2 channels have a significant electrical role, acute inhibition with GxTX-1E should significantly alter firing characteristics of both control (K_V_2.1^ON^ / K_V_2.2^ON^) and ChAT-K_V_2.1^OFF^ motoneurons (K_V_2.1^OFF^ / K_V_2.2^ON^). However, the only consistent effect we found was a relatively small (less than 10% in all neurons) reduction in the instantaneous frequency of the initial two spikes in a train. We saw the same effect in the ChAT-K_V_2.1^OFF^ motoneurons, suggesting K_V_2.2 currents also contribute to maintaining high initial firing frequencies. Functionally, this could be significant as small increases in initial firing frequencies are known to significantly increase the rate of muscle force production and its maintenance over time (Stein & Parmiggiani, 1979; Spielmann *et al*., 1993).

We note that these findings may seem to contrast with those reported by others, which indicated that K_V_2 inhibition alters motoneuron firing and excitability (Fletcher *et al*., 2017; Romer *et al*., 2019; Nascimento *et al*., 2020). Fletcher *et al*. (2017) used GxTX-1E to block motoneuron K_V_2 channels in P4 mice. They suggested that K_V_2 conductances maintain narrow spikes and repetitive firing at low current inputs, and increase firing frequencies. Romer *et al*. (2019) assessed the contribution of K_V_2 to regulation of motoneuron firing in spinal cord slices from P8-12 rats, which are approaching motor maturity. They reported that blocking K_V_2 channels with stromatoxin led to unsustainability of repetitive firing, a reduction in excitability (f-I slope), and a slowing of the rising and falling phases of action potentials (simlar to Fletcher *et al*., 2017). And Nascimento *et al*. (2020) showed in young (P2- P7) motoneurons that chemogenetic activation of V0_C_ interneurons (the neuronal source of C-boutons) decreases spike width, while increasing maximal firing rates, the current required for depolarising block, and mAHP, and these effects were all blocked by GxTX-1E. They thus suggested that clustered K_V_2.1 channels underlie these M2-mediated electrophysiological effects, and are recruited by C-boutons for motor amplification.

Our results are in keeping with the above studies showing that conduction through K_V_2 channels can affect motoneuron firing in the early post-natal period. For example, we too showed that K_V_2 channels contribute to spike width in early post-natal life, during a period when K_V_2.1 channels and motor control are immature (Altman & Sudarshan, 1975; Wilson *et al*., 2004). And the one study above in an intermediate developmental stage (Romer *et al*., 2019) used stromatoxin, which blocks K_V_2 channels, but also has off target effects (K_V_ 4.2 channels with a low IC_50_) that make interpretation of current clamp data challenging (Escoubas *et al*., 2002; Liu & Bean, 2014).

Two clear differences, in addition to the age of the preparation, between our study and that of Nascimento *et al*. (2020) are the specificity of C-bouton activation and their use of en bloc preparations. While their chemogenetic activation would limit off target effects, we used muscarine - an agonist of all muscarinic receptors that has been shown to have mixed effects on early post-natal motoneurons (Miles *et al*., 2007; Nascimento *et al*., 2019). In agreement with this, we found that in young motoneurons, muscarine led to significant depolarisations (Ghezzi *et al*., 2017). But these effects were not present beyond the 2nd post-natal week, suggesting a change in muscarine receptor effects. In these older motoneurons, the effects we report with muscarine were dominated by M2-/ C-bouton-type effects of increased excitability, and these were similar in ChAT-K_V_2.1^OFF^ motoneurons.

The use of the en bloc preparation could also lead to an increase in the basal level of C-bouton activity due to spontaneous activity in V0_C_ neurons (Zagoraiou *et al*., 2009; Nascimento *et al*., 2020), which have intersegmental connections (Stepien *et al*., 2010). Therefore, in the slice preparation, there may be insufficient basal activity to activate K_V_2.1 channels. That is, if K_V_2 channels have a significant electrical role that is only activated by C-boutons, the effects of GxTX-1E may be difficult to discern in slice preparations. However, we note that our findings are consistent with those demonstrating that highly clustered K_V_2 channels (like those opposite C-boutons) are non-conducting (O’Connell *et al*., 2010; Fox *et al*., 2013; Johnson *et al*., 2018; Maverick & Tamkun, 2022). And if that is the case, then basal V0c activity is unlikely to activate these conductances. In addition, less dense K_V_2 channels (which are more likely to be electrically active) are found opposite other (mainly excitatory) synapses (Muennich & Fyffe, 2004), which would also be more active in en bloc preparations, complicating data interpretation.

It therefore seems most likely that the differences in findings between the above studies and ours can be reconciled by considering the maturity of the preparations and the specificity of GxTX-1E for K_V_2 channels. Another possible contributing factor could be the different intracellular solutions (e.g. energy support) used in the different studies.

### Does maturation of K_V_2.1 channels influence their role in motoneurons?

The conducting state of K_V_2 channels is tightly linked to their spatial density on the plasma membrane, regardless of their state of phosphorylation (O’Connell *et al*., 2010; Fox *et al*., 2013; Maverick & Tamkun, 2022). In cells in which the majority of K_V_2.1 channels are expressed in high density domains on the PM, only a fraction are conducting and simply declustering channels via dephosphorylation does not seem to alter channel conductance (Benndorf *et al*., 1994; O’Connell *et al*., 2010; Fox *et al*., 2013; Maverick & Tamkun, 2022). Our data in motoneurons are in keeping with these findings: we show that the overall number of K_V_2.1 puncta per 100 µm^2^ of motoneuron membrane is slightly reduced with postnatal development as most channels form large, high density clusters opposed by C-boutons, and over this same period, K_V_2 channels contribute less to motoneuron electrophysiological properties. The molecular mechanisms responsible for reducing K_V_2.1 conductances in post-natal development are not known.

### K_V_2 channels play a primarily non-conducting role in motoneurons

Our muscarine and behavioural data suggested that either K_V_2.1 has a minimal conducting role in C-bouton amplification or that, because K_V_2.1 and K_V_2.2 are co-clustered (possibly heteromeric; Kihira *et al*., 2010), this role could be subsumed by K_V_2.2 alone, resulting in a safety factor in C-bouton development. Given that GxTX-1E, which blocks both K_V_2.1 and K_V_2.2 channels, had little effect on motoneuron firing, we propose that K_V_2 channels have a minimally conducting role in mature motoneurons. Of note, this finding is very different from that in other cells, such as some cortical (as above) and hippocampal pyramidal neurons (Misonou *et al*., 2004; Liu & Bean, 2014).

But there is plenty of evidence showing electrically silent voltage-gated potassium channels (for review see Bocksteins & Snyders, 2012), and we now know that K_V_2 channels have various non-conducting roles (Feinshreiber *et al*., 2010; Deutsch *et al*., 2012; Kirmiz *et al*., 2018; Vierra *et al*., 2019). In motoneurons, the paucity of effects when knocking out or blocking K_V_2.1 channels led us to shift our thinking towards the possibility that K_V_2 channels play a primarily non-conducting role in motoneurons, such as maintaining the spatial relationship between the ER and (PM, see below; Johnson *et al*., 2018), the hallmark of C-bouton synapses (Conradi, 1969). However, as the roles of subsurface cisterns and their physical links to the PM are not yet clear in motoneurons, the functional significance of activity-dependent declustering also remains obscure.

In summary, it is possible that a small fraction of motoneuron K_V_2 channels are electrically active, and these may support initial high frequency firing and higher maximal frequencies, but overall our data suggest that these channels are largely electrically silent in motoneurons.

### Possible sampling bias

Here, we targeted the largest motoneurons, reasoning that C-bouton amplification is principally evident in high motor output tasks (Zagoraiou *et al*., 2009) and is thus related to recruitment of the faster (and larger) motoneurons that innervate higher force-producing fast-twitch muscle fibres. It is possible that there are clustering/expression differences between slow and fast motoneurons, although none have yet been reported. Interestingly, neither KCNG4, a potassium channel subunit expressed in fast motoneurons (Müller *et al*., 2014), or SK3, a calcium-dependent potassium channel that may be expressed primarily in slow motoneurons (Deardorff *et al*., 2013; Kissane *et al*., 2021), has overlap in expression across motor pools with K_V_2.1 (http://spinalcordatlas.org; Blum *et al*., 2021). Nonetheless, taking the electrophysiological data together with the behavioural data, it is unlikely that a bias towards the larger motoneurons significantly impacted our conclusions.

### What role do K_V_2 channels play in C-bouton motor amplification?

To assess whether K_V_2.1 cKO influenced motor amplification, we performed whole cell current clamp experiments using muscarine to activate M2 receptors in vitro, and behavioural tasks requiring high force output to test C-bouton function in vivo. Our patch clamp experiments showed that muscarine increased excitability in ChAT-K_V_2.1^OFF^ motoneurons despite the absence of K_V_2.1 channels, suggesting K_V_2.1 is not necessary for M2R mediated motor amplification by C-boutons. This hypothesis was supported by our behavioural data as we found control and ChAT-K_V_2.1^OFF^ mice were equally capable of amplifying their motor output. Our finding that K_V_2.2 channels are co-expressed with K_V_2.1 at C-bouton synapses suggests that, if K_V_2 channels are important for neuronal function, K_V_2.2 can function independently of K_V_2.1 during behaviours requiring C-bouton activation.

What role might K_V_2 channels play at C-bouton synapses? There is strong evidence that K_V_2 channels form physical links with the ER membranes via VAP proteins in order to maintain tight PM-ER junctions (Fox *et al*., 2015; Johnson *et al*., 2018; Kirmiz *et al*., 2018; Deardorff *et al*., 2021). Because several different proteins apposing C-boutons are Ca^2+^-dependent, motoneuron subsurface cisterns and their associated Ca^2+^ stores are likely integral to function. Thus, K_V_2 channels may act to maintain the proximity of the sub-surface cisterns to the PM, such that M2 receptors can modulate local Ca^2+^ flux within the nanodomain of the C-bouton signalling ensemble.

Is there an advantage to having a voltage-dependent channel function primarily as a structural protein? Even when non-conducting, the K_V_2 voltage sensors responds to changes in membrane potential, and can be modulated by Ca^2+^ / calcineurin dependent phosphorylation (O’Connell *et al*., 2010; Maverick & Tamkun, 2022). It is plausible, therefore, that K_V_2 interactions with other proteins can be modulated in a state-dependent manner, depending on membrane voltage. For example, in cultured hippocampal neurons, K_V_2.1 promotes clustering of PM located L-type calcium channels and their functional coupling to ER ryanodine receptors, which are responsible for ER Ca^2+^ release (Vierra *et al*., 2019). While this is unlikely to be the specific case in motoneurons (given the absence of L-type calcium channels and lack of evidence of ryanodine receptors), there are many proteins at this site that could possibly be corralled by a K_V_2-dependent mechanism.

### Conclusions

Evolution led to the development of C-bouton synapses to amplify motor output in a task-dependent manner (Miles *et al*., 2007; Zagoraiou *et al*., 2009). To understand the mechanisms of this important amplification, it is necessary to identify the structure and function of the plethora of proteins clustered in this region, and to understand how these proteins interact with each other. We demonstrate that in motoneurons, post-synaptic K_V_2.1 and K_V_2.2 are co-expressed and likely serve predominantly non-conducting roles. However, the data also highlight that there is still much to learn regarding synaptic mechanisms of C-bouton motor amplification.

## Methods

### Animals

All experiments were approved by the University College London Animal Welfare and Ethical Review Body, and performed under Project Licence 70/9098 granted under the Home Office Animals (Scientific Procedures) Act 1986.

Three different mouse lines were used for this work. Wild type C57BL/6 were acquired from Charles River Laboratories, Inc (strain code:027) and used for breeding transgenic and conditional knockout (cKO) lines, and for patch clamp electrophysiology experiments targeting cortical pyramidal neurons. Heterozygous *KCNB1*^(+/lox)^ mice were acquired from the MRC Harwell facility back-crossed to C57BL/6 mice, and then bred to homozygosity (*KCNB1*^(lox/lox)^). ChAT-IRES-Cre (ChAT^(+/Cre)^) mice were acquired from JAX (B6;129S6-Chattm2(cre)Lowl/J, stock no. 006410), and maintained on a C57BL/6 background. Hb9::eGFP mice, also maintained by breeding with C57BL/6 wild-types, were used for developmental immunohistochemistry (P2-21) and patch clamp experiments (P2-7). These mice were acquired from the Jessell lab in 2000 and are now available from JAX (B6.Cg-Tg(Hlxb9-GFP)1Tmj/J, stock no. 005029).

To generate *KCNB1* cKO mice in which cholinergic neurons lack K_V_2.1 channels, *KCNB1*^(lox/lox)^ mice were crossed with ChAT^(Cre/Cre)^ mice to produce ChAT^(+/Cre)^; *KCNB1*^(+/lox)^ offspring. Cre-positive animals with a floxed allele were then crossed back to *KCNB1*^(lox/lox)^ mice to produce ChAT^(+/Cre)^; *KCNB1*^(lox/lox)^ offspring (Figure 1 - figure supplement 1).

The number and sex of animals used for each experiment are declared in the figure legends.

### Blinding and randomisation

For behavioural and anatomy experiments involving comparison of mature control and ChAT- K_V_2.1^OFF^ mice, animals were assigned to groups based on their genotype and sex, but experimenters remained blinded to group identity throughout all experiments. For electrophysiology experiments, blinding was not possible before recording, so this was done before data analysis. For developmental anatomy experiments, experimenters were blinded after imaging had taken place due to obvious anatomical differences in tissue of different ages. Sections of all ages were stained simultaneously in batches and well order was randomised.

### Immunohistochemistry

Animals were deeply anaesthetised by intraperitoneal injection of ketamine (100 mg kg^-1^) and xylazine (20 mg kg^-1^). Once insentient (loss of paw withdrawal), animals were perfused with 5 ml of phosphate buffered saline (PBS) followed by 20 ml of 4% paraformaldehyde. Vertebral columns were dissected and post-fixed for 24 hours before spinal cords were dissected and cryoprotected in 30% sucrose for 72 hours. Then, meninges were removed, and regions of interest segmented, frozen in OCT solution, and stored at −20 ℃. Transverse sections of 30-50 µm were cut using a cryostat (CM3050 S, Leica) and stored in 1x PBS as free floating sections.

To visualise motoneurons, C-boutons, and K_V_2.1 channels, sections were washed 3 times in PBS (10 minutes per wash) and then incubated for 1 hour in blocking solution containing 3% normal donkey serum (NDS, SC30-100ML, Merk Millipore) and 0.2% Triton X-100 (648464, Merk Millipore) diluted in PBS (PBST). Sections were then incubated in blocking solution containing goat anti-choline acetyltransferase (ChAT, 1:250, Millipore Cat# AB144P, RRID:AB_2079751) and rabbit anti- K_V_2.1 (1:500, Millipore Cat# AB5186-200UL, RRID:AB_2131651) antibodies for 48 hours at 4℃, then washed (3 x 10 minutes in 1xPBS) and incubated for 2 hours at room temperature (22 ℃) in blocking solution containing the secondary antibodies Alexa Fluor® 555 donkey anti-rabbit (1:500, Thermo Fisher Scientific Cat# A-31570, RRID: AB_2563181,) and Alexa Fluor® 488 donkey anti-goat (Thermo Fisher Scientific Cat# A-11055, RRID: AB_2534102, 1:500). Finally, sections were washed (3 x 10 minutes in 1xPBS) before being mounted on glass slides with Mowoil 4-88 (Carl Roth GmbH & Co. Kg).

To visualise C-boutons and K_V_2.2 channels, staining for goat anti-ChAT and mouse anti- K_V_2.2 IgG1 (UC Davis/NIH NeuroMab Facility, K37/89, RRID:AB_2750662, 1:200) were done sequentially (ChAT first then K_V_2.2) using the same incubation times as above for each step. The same primary, secondary, and blocking solutions and concentrations were used for ChAT staining. For K_V_2.2 channels, NDS was substituted for normal goat serum (NGS, 3%) and Alexa Fluor® 555 goat anti-mouse IgG1 (Thermo Fisher Scientific Cat# A-21127, RRID:AB_141596, 1:500) was the secondary antibody.

To determine whether the knockout strategy was successful, control (females, animals=3, slices=3 per animal, age=P21) and cKO (females, animals=3, Slices=3 per animal, age=P21) slices stained for ChAT and K_V_2.1 were imaged using a Zeiss LSM 800 confocal microscope (Zeiss LSM 800 inverted confocal microscope with Airyscan, RRID:SCR_015963), 20× objective (1 AU aperture) and Zeiss ZEN Blue Edition software (ZEN Digital Imaging for Light Microscopy, RRID:SCR_013672). Tile scan images of spinal cord hemi-sections were stitched and opened in Image J analysis software (RRID:SCR_002285) for processing. Using the particle analysis package, thresholding was performed on all images using the same settings to produce a black and white image showing only K_V_2.1 puncta. Regions of interest (ROI) of the same dimensions were defined to segment analyses of dorsal, intermediate, and ventral laminae. The nucleus counter function was then used to automatically produce a count of all puncta within the ROI, which was expressed as density (number per 100 µm^2^).

#### Postnatal development of K_V_2.1 and C-boutons on motoneurons

Procedures for assessing the postnatal development of C-boutons and K_V_2.1 channels on lumbar motoneurons were largely the same as those described above, with a few differences. Lumbar (L4-5) sections were cut (50 µm) from neonatal (P2-3), transition (P6-7) and motor mature (P21) Hb9::eGFP transgenic mouse spinal cords. Vesicular Acetylcholine Transferase polyclonal antibody (VAChT, 1:500, Millipore Cat# ABN100, RRID:AB_2630394) was used to visualize C-boutons and K_V_2.1 was visualised using the same antibody as described above. Secondary antibodies used were Alexa Fluor® 647 donkey anti-goat (1:500, Thermo Fisher Scientific Cat# A-21447, AB_2535864) and Alexa Fluor® 555 donkey anti-rabbit (1:500, Thermo Fisher Scientific Cat# A-31570, RRID: AB_2563181).

60x confocal z-stack (0.4µm steps through tissue thickness) images were captured with the LSM 800 inverted confocal microscope and then pseudo-named in order to perform blinded analyses. Three-dimensional (3D) reconstructions of each motoneuron were then rendered using IMARIS Software (Bitplane, RRID:SCR_007370) using the following procedure. First, in the 3D isometric view, solid surfaces of the motoneuron soma (Hb9::eGFP signal), C-boutons, and K_V_2.1 were created using the rendering and thresholding tools. The masking feature was then used to select K_V_2.1 clusters within 1 µm of the motoneuron surface and VAChT^+^ C-boutons. IMARIS was used to generate volume and surface area data for each motoneuron and associated C-boutons and K_V_2.1 clusters, and these were exported to an Excel (Microsoft Corporation, 2018, RRID:SCR_016137) spreadsheet. Subsequent analyses were performed using python programming language run in Jupyter Labs environment. Cells were excluded only if quality of staining precluded accurate rendering of the cell.

Intensity plots were made in ImageJ by drawing a ROI with a polygon line around the perimeter of the cell, to connecting the centre points of all C-bouton puncta. The same ROI was copied to the K_V_2.1 channel and the relative intensities were measured (intensity/maximum intensity) for C-boutons and K_V_2.1. The relative intensity values (y-axis) were then plotted against the corresponding distance values (x-axis).

### Patch clamp electrophysiology

#### Slice preparation

Patch clamp electrophysiology experiments were performed as described inSmith and Brownstone (2020). Mice of all ages were administered an intraperitoneal bolus of ketamine (100 mg kg-1) and xylazine (20 mg kg-1) and decapitated following loss of hind-paw withdrawal. The vertebral column was quickly excised and pinned (ventral-side-up) to a silicone dish containing ice-cold (0–4℃) normal artificial cerebrospinal fluid (nACSF) saturated with 95% carbogen. nACSF was made in 18 MΩ water with the following (in mM): 113 NaCl, 3 KCL, 25 NaHCO3, 1NaH2PO4, 2 CaCl, 2 MgCl2 and 11 D-glucose, pH 7.4 (Mitra & Brownstone, 2012). A vertebrectomy was performed to reveal the spinal cord, which was stripped of dura matter and extracted from the vertebral column.

The spinal cord was then glued (3M Vetbond, No.1469SB) ventral-side-up to a pre-cut block of agarose (∼8% in ddH_2_O) and mounted on a cutting chuck using superglue. These steps were completed as quickly as possible to ensure viability of slices from animals in the third post-natal week, typically within 3 minutes.

The chuck was transferred to the slicing chamber of a vibrating microtome (Model 7000 smz-2, Campden Instruments Ltd) containing ice-cold slicing solution made up of the following (in mM): 130 potassium gluconate, 15 KCL, 0.05 EGTA, 20 HEPES, 25 glucose, 3 kynurenic acid, pH 7.4 (Dugué *et al*., 2005; Bhumbra & Beato, 2018). 350 µm slices were cut and transferred to the incubation chamber to rest in nACSF (32℃) for 30 minutes. The incubation chamber was then allowed to equilibrate to room temperature (maintained at 23°C) for at least 30 minutes before recording.

#### Recording and analyses

Using a DMLFSA microscope (Leica DMLFSA; Leica Microsystems,Wetzlar, Germany), putative motoneurons were identified as the largest cells in the motor pools of the spinal cord. Patch pipettes were pulled using a P97 Flaming/Brown horizontal Micropipette Puller (Sutter Instrument, RRID:SCR_016842) to a resistance of 1.5-4 MΩ. Patch pipette electrodes were filled with an internal solution consisting of (in mM): 131 K-methanesulfonate, 6 NaCl, 0.1 CaCl2, 1.1 EGTA, 10 HEPES, 0.3 MgCl2, 3 ATP-Mg, 0.5 GTP-Na, 2.5 L glutathionine, 5 phosphocreatine, pH 7.25 adjusted with KOH, osmolarity 290–300 mOsm.

Recordings were made using a MultiClamp 700A amplifier (Axon Instruments, Inc), low pass filtered at 10 kHz and digitized at 25 kHz using a CED Power3 1401 (Cambridge Electronic Designs Limited). All experiments were performed in current clamp mode and data captured using Signal software (Cambridge Electronic Design Ltd, Cambridge, UK, RRID:SCR_017282). Once whole cell configuration was achieved, the bridge was balanced, and capacitance neutralized prior to commencing recording. Motoneurons were injected with a small negative rectangular pulse (500 ms duration) and the voltage responses of 15–30 traces were averaged to measure input resistance and whole-cell capacitance (WCC). Resistance was measured as the peak voltage change to the injected current and τ calculated from an exponential curve fitted to the response (automated in Signal). WCC was calculated using resistance and τ values and cross-checked against the values automatically recorded by the software during the experiment. Rheobase was defined as the minimum amount of current needed to evoke an action potential. Motoneuron frequency-current (ƒ-I) graphs were generated by injecting depolarizing current steps increasing from 0 nA until maximum firing was observed. The excitability of the cell (gain) was determined by measuring the slope of the main linear portion of ƒ-I plots for first interval frequency (instantaneous frequency of the first two spikes), and overall frequency (mean instantaneous frequency of all spike in a train). Action potential half widths (1/2 width), spike amplitude, and fast afterhyperpolarization (fAHP) were measured from 15–30 averaged single APs evoked with a 20 ms rectangular current pulse (to ensure stimulus artefact did not preclude measurement). The ½ width was calculated as the time between the 50% rise and 50% fall in amplitude of the AP. Spike height was measured as the voltage difference between the threshold (voltage at maximum positive value of the second derivative of membrane potential of AP) and the peak of the AP. The fAHP was measured as the difference between the voltage baseline and the most negative point on the first trough of the AP. Afterpotential measurements (mAHP amplitude and mAHP half decay time) were taken from averages of 15-30 single APs evoked with a 1 ms duration current pulse (to ensure stimulus artefact preceded mAHP). The mAHP amplitude was calculated from baseline to the most negative point on the trough. The mAHP half decay time is calculated as half the time taken (ms) from the most negative point of the mAHP to baseline. Cells were held at −65 mV for single evoked APs.

#### Mature motoneurons

For these experiments, the experimental unit (N) was considered to be individual motoneurons and is disclosed in the results section and figure legend (as well as animal number and sex) for each experiment. To avoid sampling γ-motoneurons we selected the largest motorneurons in motor pools with resting membrane properties consistent with those of verified ɑ-motoneurons-this was determined *a priori*. As a result, the majority of motoneurons sampled had input resistance values ≤ 50 MΩ. During analysis, we discovered a sampling bias between control and cKO motoneurons, whereby the control group had disproportionately more neurons with input resistance values > 50 MΩ. To account for this we only included motoneurons with input resistance values ≤ 50 MΩ (Figure 2 - figure supplement 1B).

#### Drugs and toxins

Baseline firing characteristics were recorded in nACSF for all experiments prior to the perfusion of drugs or toxins. To assess the effect of inhibiting K_V_2 channels, 100 nM GxTX-1E (Tocris, cat# 5676) in nACSF was perfused through the recording chamber for 10 minutes prior to the 2^nd^ recording (Liu & Bean, 2014; Fletcher *et al*., 2017; Newkirk *et al*., 2022). For experiments using muscarine (10μM, Sigma-Aldrich, cat# M6532), slices were perfused for 5 minutes prior to the 2^nd^ recording (Miles *et al*., 2007; Ghezzi *et al*., 2017; Nascimento *et al*., 2019).

#### Cortical pyramidal neuron patch clamp experiments

Coronal brain slices (350 μm) were made from wild type C57BL/6 male mice aged P14-15 (N=3). Whole cell, current clamp recordings were made from visually identified pyramidal neurons in layer 5 of the cerebral cortex. Subsequent procedures were performed as described for mature motoneurons above.

### EMG recording for C-bouton motor amplification

Bipolar, intramuscular electrodes were fabricated using materials and methods described in detail by Pearson *et al*. (2005).

Procedures for surgical implantation were also based on those first described by Pearson *et al*. (2005). Mice were anaesthetised with isoflurane (5% with O_2_) and tested for loss of the paw withdrawal reflex before proceeding. Once insentient, the back and both hindlimbs were shaved, cleaned with 70% ethanol and then surgical iodine solution. An incision measuring the width of the EMG connector was made between the scapulae and another of similar size was made in the centre of the back at the level of the hips. Wires from the connector were tunnelled from the rostral to the caudal incision and the connector was secured in place using 4-0 sutures (ETHICON, W8683). Incisions were then made over the medial gastrocnemius and tibialis anterior of each hindlimb and corresponding wires tunnelled from the rostral incision to the muscle. Wires were inserted through the belly of the muscle up to the pre-made proximal knot in the wires and another distal knot was made to secure the recording sites in to muscle. The wires were trimmed, all incisions were washed with saline, and then closed with 7-0 sutures (ETHICON, W8702). Prior to withdrawing anaesthesia the mouse was administered buprenorphine (Vetergesic, 0.1mg/kg) and then transferred to a recover chamber maintained at 37 ℃ until fully awake and ambulatory. Mice were given 1 drop of oral Metacam (Meloxicam, 1.5 mg/ml) analgesic daily for 5 days post-surgery, or as long as necessary.

After at least 10 full days of recovery, EMG recording sessions were undertaken. Immediately prior to recording, mice were lightly anaesthetised with isoflurane and the male connector was inserted into the female connector (sutured into the skin at the back of the neck). Once awake and ambulatory, mice were placed on a treadmill set to a speed of 0.15 m/s and recording commenced. Following walking experiments animals were transferred to a custom swimming pool (25 ℃) and muscle activity during swimming was recorded. Signals were amplified (x 1000) using an NL844 AC pre-amplifier (Digitimer Ltd) connected to an NL820 isolation amplifier (Digitimer Ltd), filtered (100 Hz-10KHz) and digitized with a Power 1401 interface and Spike2 software (Cambridge Electronic Design, Cambridge, United Kingdom).

### Behavioural assessments and analysis

#### Running wheel experiments

Experimental mice were housed in large (rat) cages (Allentown, NexGen Rat 900) separated into 2 compartments with a perforated Perspex divider. One compartment housed a companion mouse, that had no access to a running wheel and the other was occupied by either a control (ChAT^(wt/wt)^;KCNB1^(f/f)^) or KO mouse (ChAT^(Cre/wt)^;KCNB1^(f/f)^) which had *ad libitum* access to an externally mounted running wheel (Panlab, LE905). Mice underwent an initial acclimation period (7 days) in which they had no access to the running wheel. Following acclimation, mice had 24 hour access to the wheels and data were collected (Panlab multicounter, LE3806) in either 5 or 1 minute epochs for a total of 16 hours from 19:00 to 11:00. Running distances were calculated from the circumference of the wheels (50.24 cm) and the number of revolutions. Experiments lasted 5 weeks in males and 6 weeks in females, after which data were exported to Microsoft Excel spreadsheets and analysed using Python scripts in the Jupyter notebooks environment. Animals were excluded only if determined to be non-compliant. Non-compliance was determined as the mouse running less than 500 metres per 16 hour period. Only 1 mouse was excluded for non-compliance.

#### Treadmill speed and endurance experiments

Mice were acclimatised to the treadmill (Panlab, multilane treadmill, LE8710MTS) for a period of 3 days before testing: On day 1, the belt was kept static and mice were allowed to explore for 10 minutes. On day 2, the treadmill was set to a 15% incline and mice walked slowly (5 cm/s) for a total time of 15 minutes. The air puff encouragement was activated when contact was made with a grid at the rear of the treadmill. Day 3 consisted of a 20 minute session consisting of 2 minutes at 5 cm/s followed by increments of 2 cm/s every 2 minutes. On maximum speed testing days, mice were placed in the treadmill and the speed increased by 3 cm/s every minute for 5 minutes to warm up. Then, the speed was increased by 5 cm/s every 20 seconds until mice were persistently lagging (i.e. when total air puff stimulation reached 10s). After 2 days recovery, mice were subjected to endurance tests involving the same 5 minute warm up followed by up to 40 minutes (time cap) running at 60% of the maximum speed attained by individual mice. Again, the limit was determined as the time/distance at which mice received 10 seconds of air of stimulation.

#### Grip strength experiments

Grip strength was tested using a BIOSEB grip strength meter (Figure 9- figure supplement 3) over 3 days, with 2 days rest between each session. The experimenter held the tail of the mouse whilst supporting its weight on their other hand before lowering it onto the horizontal metal grid attached to the grip strength meter. Once the mouse gripped the bars, the experimenter slowly pulled backwards by the tail until the mouse released its grip. The peak force (grams) was recorded and the mouse was returned to its cage to rest for 1 minute. This process was repeated 3 times for each mouse on each day of testing. Thus, mean force outputs (normalised to weight g/g body weight) for each animal were calculated from a total of 9 ‘pulls’.

### Statistical methods

We centre our analyses on estimation statistics because it focusses conclusions on magnitude, precision and biological significance of the results, thereby circumventing many of the flaws associated with null hypothesis significance testing (NHST; Bernard, 2019; Makin & de Xivry, 2019; Wasserstein *et al*., 2019; Michel *et al*., 2020). Hedges g effect sizes are calculated along with bootstrapped confidence intervals, which describe the range of effect sizes possible, rather than providing a single dichotomous decision (as done in NHST). Effect sizes can be classified as no effect (0-0.19), small (0.2-0.49), medium (0.5-0.79) and large (0.8; Hedges, 1981). Boostrapped confidence intervals (CI) are used to determine the precision of the effect size; for confidence intervals that do not include 0, effect sizes are considered precise enough to attribute biological significance to the observed effects.

For unpaired comparisons of two groups, Gardner-Altman estimation plots were used (Altman *et al*., 2013), where the groups are plotted on the left axes and the Hedges g effect size with bootstrap resampled (5000 reshuffles) 95% confidence intervals plotted on the right. The Hedges g effect size is depicted as a dot; the 95% confidence interval is indicated by the vertical error bar. The values for mean difference (μ), effect size (es) and 95% confidence intervals [lower, upper] are displayed each plot. Experimental units (N) are described in the figure legends.

For paired experiments, Cumming paired estimation plots were used where the experimental units are plotted on the upper graphs and each paired set of observations is connected by a line. On the lower plots, effect sizes (Hedges g) are plotted with bootstrap resampled (5000 reshuffles) 95% confidence intervals. Effect sizes are depicted as dots; 95% confidence intervals are indicated by the vertical error bars. The bootstrapped mean differences (μ) are shown on the upper plots, and the effect sizes (es) and 95% confidence intervals [lower, upper] are displayed on the lower plots.

For developmental anatomy data, Spearman’s rank correlation analyses were used to determine if there was a significant relationship between age and dependent variables. For IHC experiments assessing C-bouton and K_V_2.1 channel expression and clustering during development, 3 animals (male) were used for each age group from which 3 sections and 16-18 motoneurons (maximum 6 per slice) were sampled. Because we were able to sample the entire motoneuron soma for puncta density measurements, we consider the motoneuron to be the experimental unit in these experiments.

For analysis of EMG data, signals acquired during walking and swimming were converted to RMS amplitudes (τ = 40 ms) and the peak values of each burst for each muscle were averaged. Motor amplification ratio was expressed as mean RMS amplitude during swimming / walking. Mean amplitudes across muscles (MG or TA) were treated as individual experimental units. BCa bootstrap resampling and permutation t-tests were used as described above for unpaired groups. For wheel running experiments assessing distance run per day, repeated measured ANOVAs were used as a suitable alternative was not available for estimation statistics.

All figures were created using Seaborn v.0.10.0 (Waskom *et al*., 2020), DABEST (Ho *et al*., 2019) and Matplotlib v.3.1.3 (Hunter, 2007) in the Python environment.

## Supporting information

Smith Brownstone supplementary figures

## Data availability

All data and Python analysis notebooks have been deposited in github. https://github.com/Brownstone-lab/KV2_paper_Allfiles_for_jupyterNB

## Materials availability statement

Newly created mice with floxed kncwill be made available upon reasonable request.

## Acknowledgements

We thank Shireen Salem for help with staining and imaging of spinal cord slices for development experiments, Nadine Simons-Weidenmaier for technical assistance and managing mouse colonies, and Filipe Nascimento and Marco Beato for helpful comments on the manuscript.

