## Supplementary material for "Kv2 conductances are not required for C-bouton mediated enhancement of motoneuron output": Smith Brownstone supplementary figures

Supplemental Figures

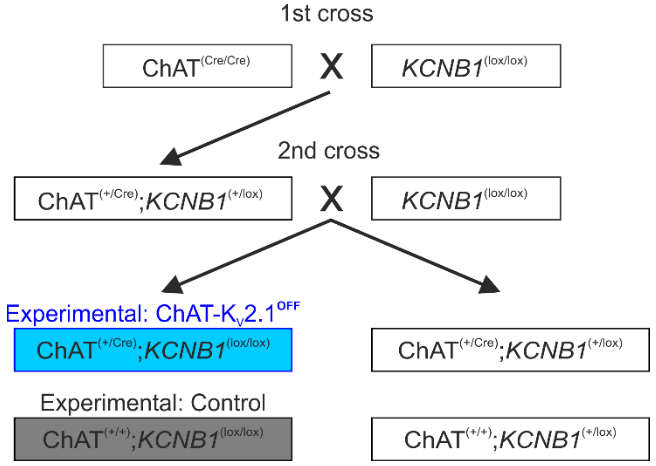


**Figure 1- figure supplement 1. Breeding strategy for conditional knockout of $K_V$2.1 in ChAT$^+$ lumbar motoneurons.** Two main breeding steps were used to generate experimental animals. In step 1 (1$^{st}$ cross), homozygous ChAT$^{(cre/cre)}$ mice were crossed with *KNCB1*$^{(lox/lox)}$ mice to produce heterozygous ChAT$^{(+/cre)}$; *KNCB1*$^{(+/lox)}$ offspring. For step 2, ChAT$^{(+/cre)}$; *KNCB1*$^{(+/lox)}$ mice were bred with *KNCB1*$^{(lox/lox)}$ mice to produce experimental (blue) ChAT$^{(+/cre)}$; *KNCB1*$^{(lox/lox)}$ mice and control mice (grey).

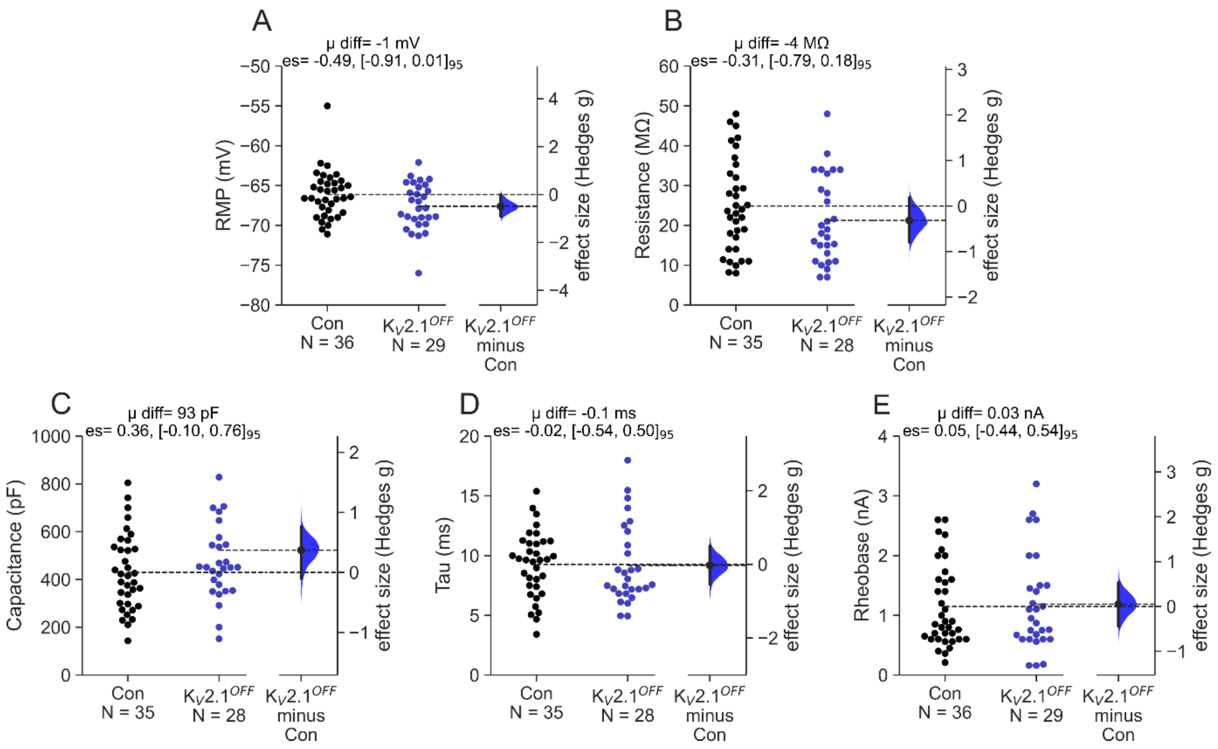


**Figure 2- figure supplement 1. $K_V$2.1 cKO does not alter motoneuron passive membrane properties. (A-E)** Gardner-Altman estimation plots. Control and ChAT-$K_V2.1^{OFF}$ groups are plotted on the left axes and the bootstrapped sampling distribution (5000 reshuffles) for Hedges g effect size are plotted on the right for resting membrane potential (RMP, A), input resistance (B), whole cell capacitance (C), time constant (tau, D), and rheobase (E). The Hedges g effect size is depicted as a dot, with 95% confidence intervals indicated by vertical error bars. The mean difference (μ diff), effect sizes (es) and 95% confidence intervals [lower, upper] are displayed at the top of each plot. Experimental unit (N) = motoneurons recorded from 23 control (n= 9 females, 14 males) and 14 ChAT-$K_V2.1^{OFF}$ mice (n= 7 females, 7 males).

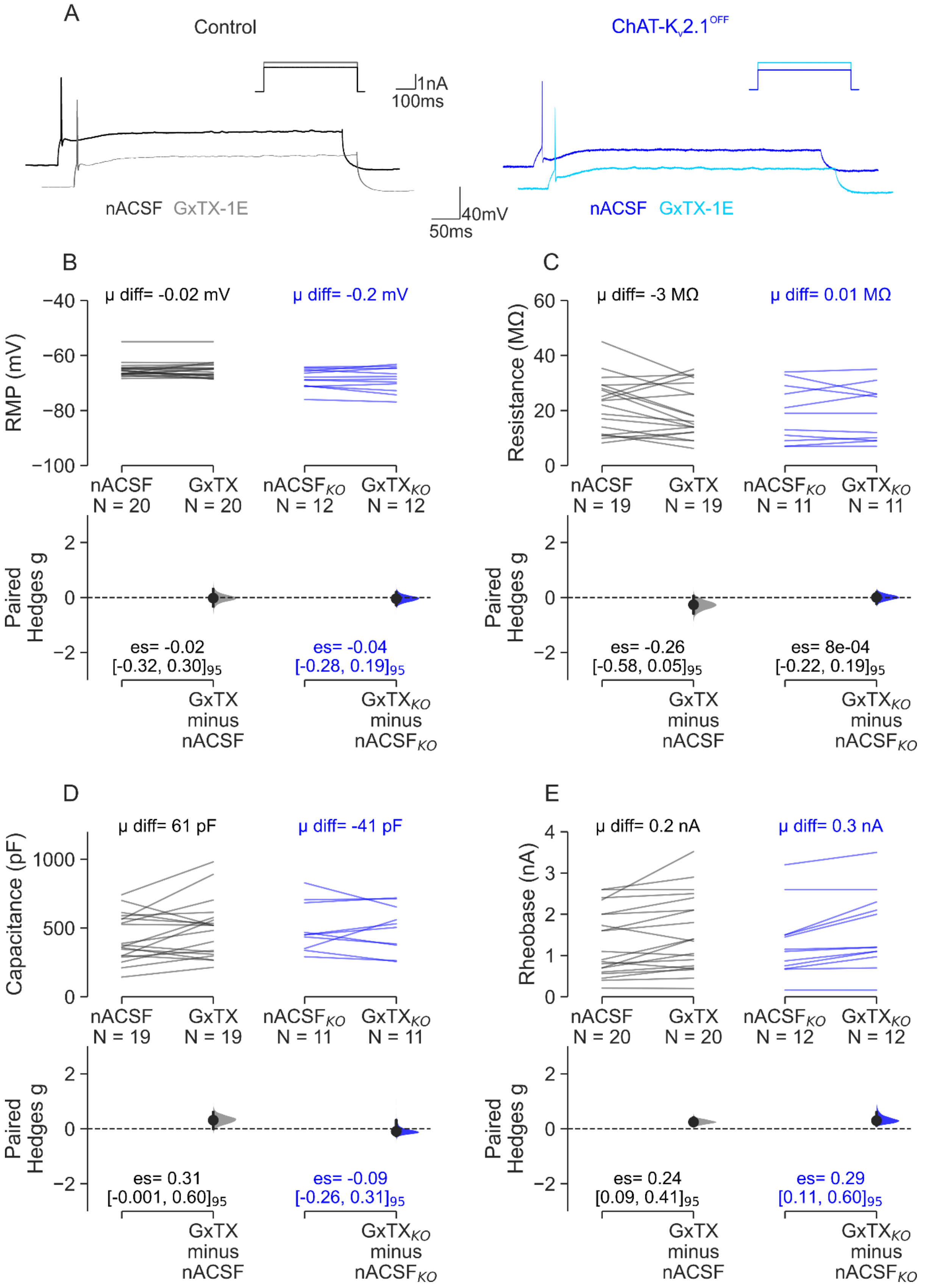
A
Control
ChAT-Kv2.1OFF
1nA
100ms
40mV
50ms
nACSF GxTX-1E
nACSF GxTX-1E
B
μ diff= -0.02 mV
μ diff= -0.2 mV
RMP (mV)
nACSF N = 20
GxTX N = 20
nACSFKO N = 12
GxTXKO N = 12
Paired Hedges g
es= -0.02 [-0.32, 0.30]95
es= -0.04 [-0.28, 0.19]95
GxTX minus nACSF
GxTXKO minus nACSFKO
C
μ diff= -3 MΩ
μ diff= 0.01 MΩ
Resistance (MΩ)
nACSF N = 19
GxTX N = 19
nACSFKO N = 11
GxTXKO N = 11
es= -0.26 [-0.58, 0.05]95
es= 8e-04 [-0.22, 0.19]95
D
μ diff= 61 pF
μ diff= -41 pF
Capacitance (pF)
nACSF N = 19
GxTX N = 19
nACSFKO N = 11
GxTXKO N = 11
es= 0.31 [-0.001, 0.60]95
es= -0.09 [-0.26, 0.31]95
E
μ diff= 0.2 nA
μ diff= 0.3 nA
Rheobase (nA)
nACSF N = 20
GxTX N = 20
nACSFKO N = 12
GxTXKO N = 12
es= 0.24 [0.09, 0.41]95
es= 0.29 [0.11, 0.60]95

**Figure 4- figure supplement 1. The effect of GxTX-1E on passive membrane properties. (A-B)** Representative traces from control **(A)** and ChAT-$K_V$2.1$^{\mathbf{OFF}}$ **(B)** motoneurons depicting a slight increase in rheobase following perfusion with 100 nM GxTX-1E. **(C-F)** Paired Hedges g for control (left) and ChAT-$K_V$2.1$^{OFF}$ (right) motoneurons in Cumming paired estimation plots, showing resting membrane potential **(RMP, C)**, input resistance **(D)**, whole cell capacitance **(E)**, and rheobase changes in response to toxin **(F)**. Individual motoneurons are plotted on the upper graphs, with each paired set of observations (nACSF followed by 100 nM GxTX-1E) connected by a line. On the lower plots, effect sizes (Hedges g) are plotted with bootstrapped sampling distribution (5000 reshuffles). Effect sizes are depicted as dots; 95% confidence intervals are indicated by the vertical error bars. The bootstrapped mean differences (μ diff) are shown on the upper plots, and the effect sizes (es) and 95% confidence intervals [lower, upper] are displayed on the lower plots. Experimental unit (N) = motoneurons. Number of animals used was as follows: control= 13 (6 females, 7 males), ChAT-$K_V$2.1$^{\mathbf{OFF}}$ = 6 (4 females, 2 males).

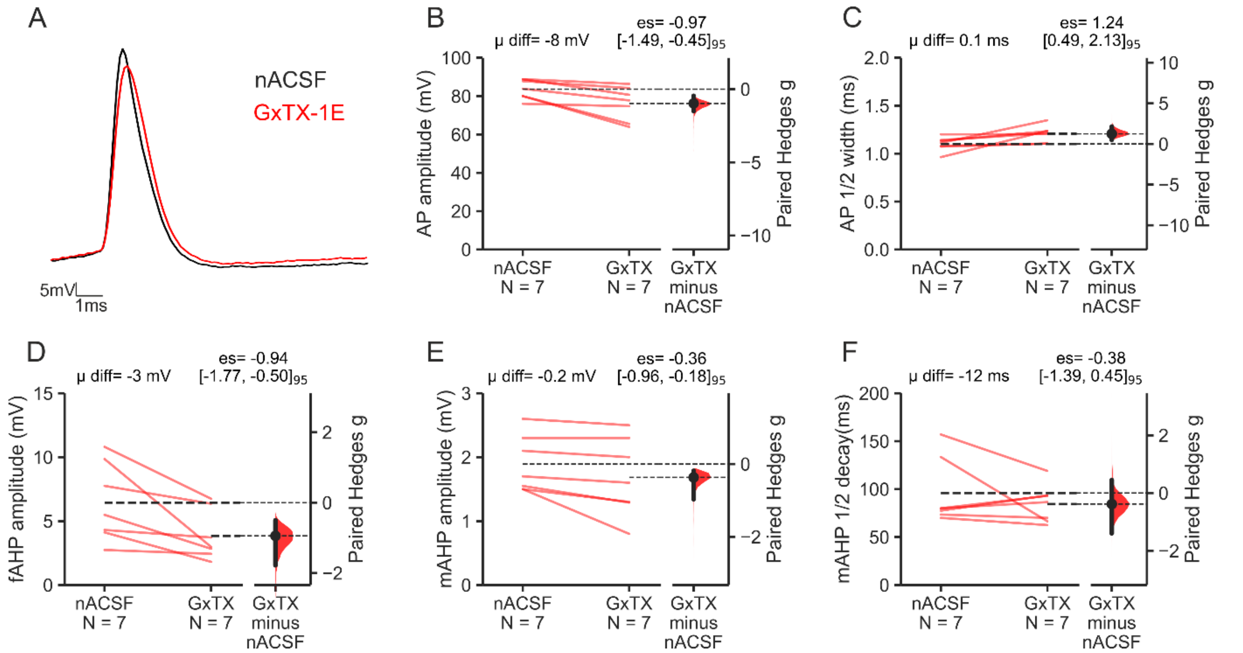


**Figure 6- figure supplement 1. Comparing the effects of $K_V2$ inhibition with GxTX-1E on cortex pyramidal neuron action potential characteristics in control mice. (A)** Representative single AP traces from layer V cortical pyradmidal neurons showing the difference in morphology pre (nACSF, black) and post 10 minute perfusion with 100 nM GxTX-1E (red). **(B-F)** Paired Hedges g for wild-type cortical pyramidal neurons are shown in Cumming paired estimation plots. Individual neurons are plotted on the upper plots, with each paired set of observations (nACSF followed by 100 nM GxTX-1E) connected by a line. On the lower plots, effect sizes (Hedges g) are shown as a bootstrapped sampling distribution (5000 reshuffles). Effect sizes are depicted as dots; 95% confidence intervals are indicated by the vertical error bars. The bootstrapped mean differences (μ diff) are shown on the upper plots, and the effect sizes (es) and 95% confidence intervals [lower, upper] are displayed on the lower plots. **(B)** shows the spike amplitude, **(C)** is action potential ½ width, **(D)** is the fAHP amplitude, **(E)** is the mAHP amplitude, and **(F)** is the mAHP ½ decay time. Experimental unit (N) = neurons recorded from 3 animals (2 males, 1 female).

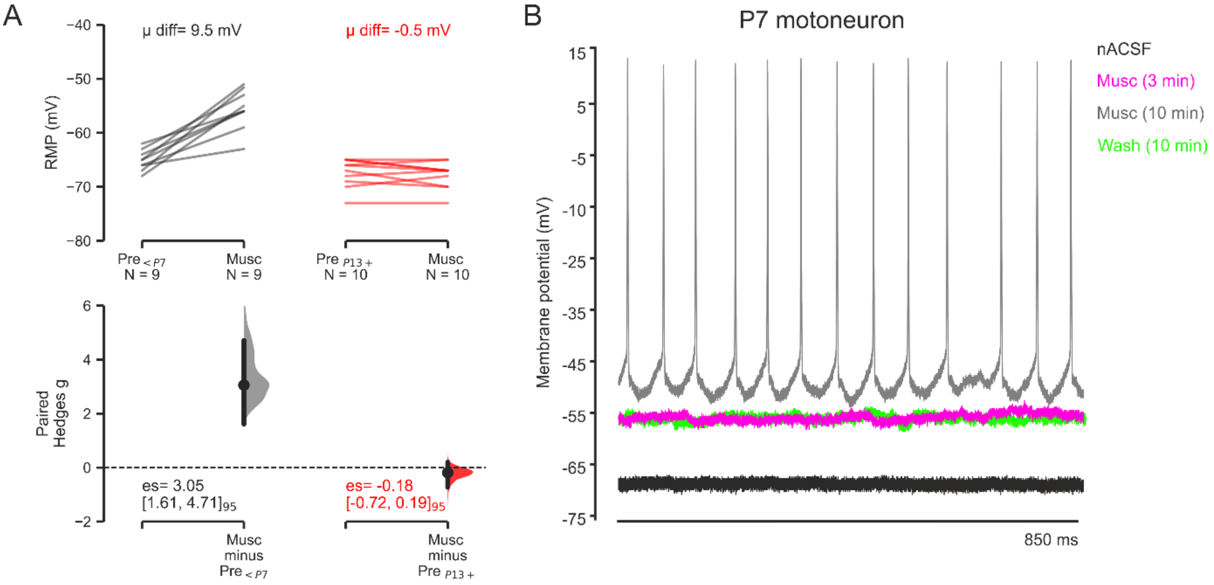


**Figure 8- figure supplement 1. Muscarine induced depolarisations in P2-7 motoneurons are abolished by P13. (A)** Paired Hedges g for young (P2-P7, left) and mature (P13+, right) motoneurons are shown in Cumming paired estimation plots. Individual motoneurons are plotted on the upper graphs; each paired set of observations (nACSF followed by 10 µM Muscarine) connected by a line. On the lower plots, effect sizes (Hedges g) are plotted as a bootstrapped sampling distribution (5000 reshuffles). Effect sizes are depicted as dots; 95% confidence intervals are indicated by the vertical error bars. The bootstrapped mean differences (µ diff) are shown on the upper plots, and the effect sizes (es) and 95% confidence intervals [lower, upper] are displayed on the lower plots. **(B)** Traces from a P7 motoneuron showing that 10 µM muscarine induces large depolarisations, and sometimes AP firing in young motoneurons. No significant depolarisations were reported in motoneurons from animals older that P13. Animal numbers were as follows: P2-3 = 6 female (8 MNs), 1 male (2 MNs); P13-21 = 4 females (8 MNs), 2 males (2 MNs).

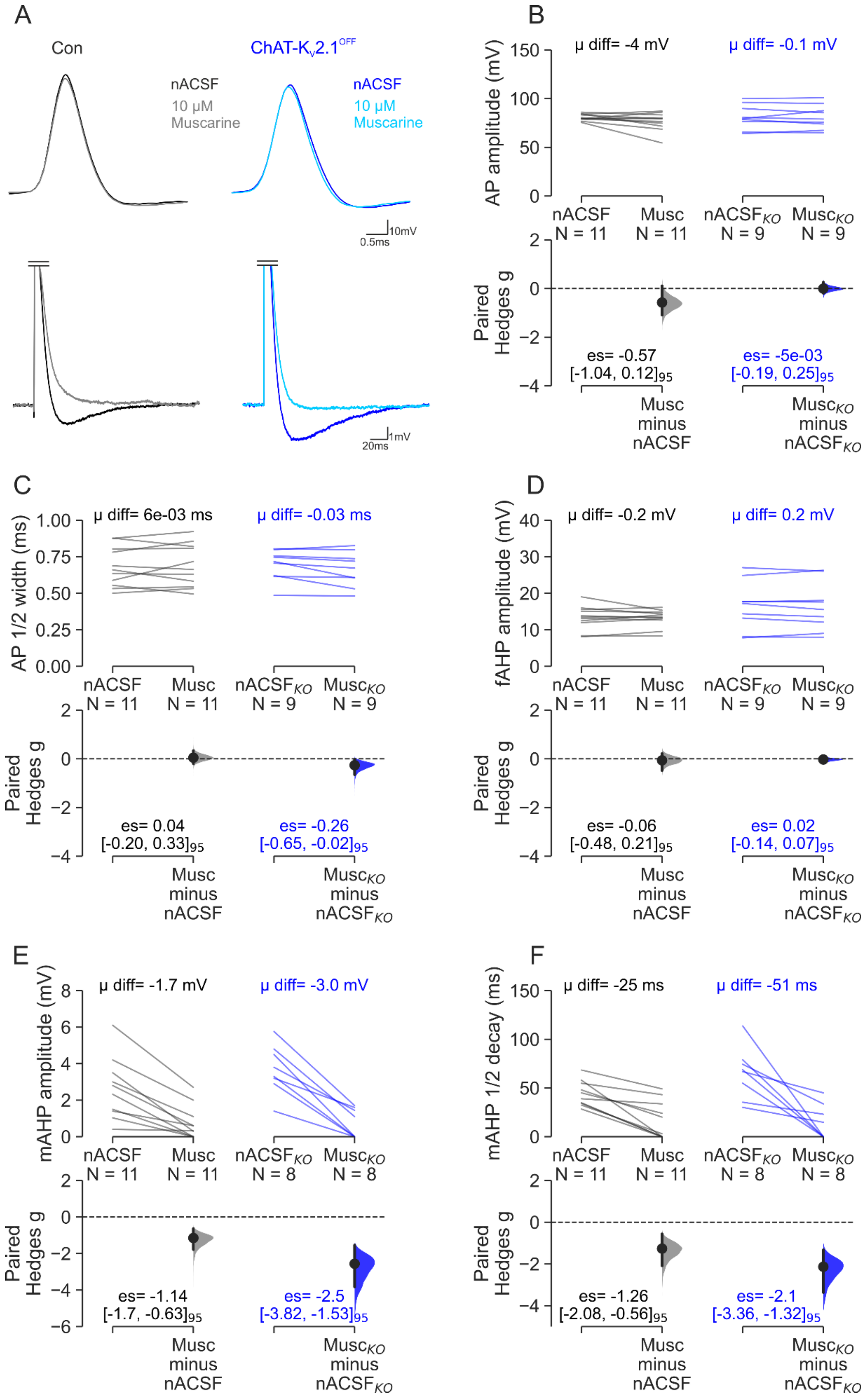

A
Con
nACSF
10 μM Muscarine
ChAT-Kv2.1OFF
nACSF
10 μM Muscarine
10mV
0.5ms
1mV
20ms
B
AP amplitude (mV)
μ diff= -4 mV
μ diff= -0.1 mV
nACSF N = 11
Musc N = 11
nACSFKO N = 9
MuscKO N = 9
Paired Hedges g
es= -0.57 [-1.04, 0.12]95
es= -5e-03 [-0.19, 0.25]95
Musc minus nACSF
MuscKO minus nACSFKO
C
AP 1/2 width (ms)
μ diff= 6e-03 ms
μ diff= -0.03 ms
es= 0.04 [-0.20, 0.33]95
es= -0.26 [-0.65, -0.02]95
D
fAHP amplitude (mV)
μ diff= -0.2 mV
μ diff= 0.2 mV
es= -0.06 [-0.48, 0.21]95
es= 0.02 [-0.14, 0.07]95
E
mAHP amplitude (mV)
μ diff= -1.7 mV
μ diff= -3.0 mV
nACSFKO N = 8
MuscKO N = 8
es= -1.14 [-1.7, -0.63]95
es= -2.5 [-3.82, -1.53]95
F
mAHP 1/2 decay (ms)
μ diff= -25 ms
μ diff= -51 ms
es= -1.26 [-2.08, -0.56]95
es= -2.1 [-3.36, -1.32]95

**Figure 8- figure supplement 2. Muscarine decreases mAHP in both control and ChAT-$K_V$2.1$^{OFF}$ motoneurons. (A)** Representative traces showing the effect of 10 µM muscarine on action potential spike (upper panels) and mAHP morphology (lower panels) in control (left) and ChAT-$K_V$2.1$^{OFF}$ motoneurons (right). Double line shows truncation for visual purposes. **(B-F)** Paired Hedges g for control (left) and ChAT-$K_V$2.1$^{OFF}$ (right) motoneurons are shown in Cumming paired estimation plots. Individual motoneurons are plotted on the upper graphs; each paired set of observations (nACSF followed by 10 µM Muscarine) connected by a line. On the lower plots, effect sizes (Hedges g) are plotted as a bootstrapped sampling distribution (5000 reshuffles). Effect sizes are depicted as dots; 95% confidence intervals are indicated by the vertical error bars. The bootstrapped mean differences (µ diff) are shown on the upper plots, and the effect sizes (es) and 95% confidence intervals [lower, upper] are displayed on the lower plots. **(B)** shows the spike amplitude, **(C)** is action potential ½ width, **(D)** the fAHP amplitude, **(E)** is the mAHP amplitude, and **(F)** is the mAHP ½ decay time. Experimental unit (N) = motoneurons. Number of animals used was as follows: control= 8 (2 females, 6 males), ChAT-$K_V$2.1$^{\mathbf{OFF}}$ = 6 (2 females, 4 males).

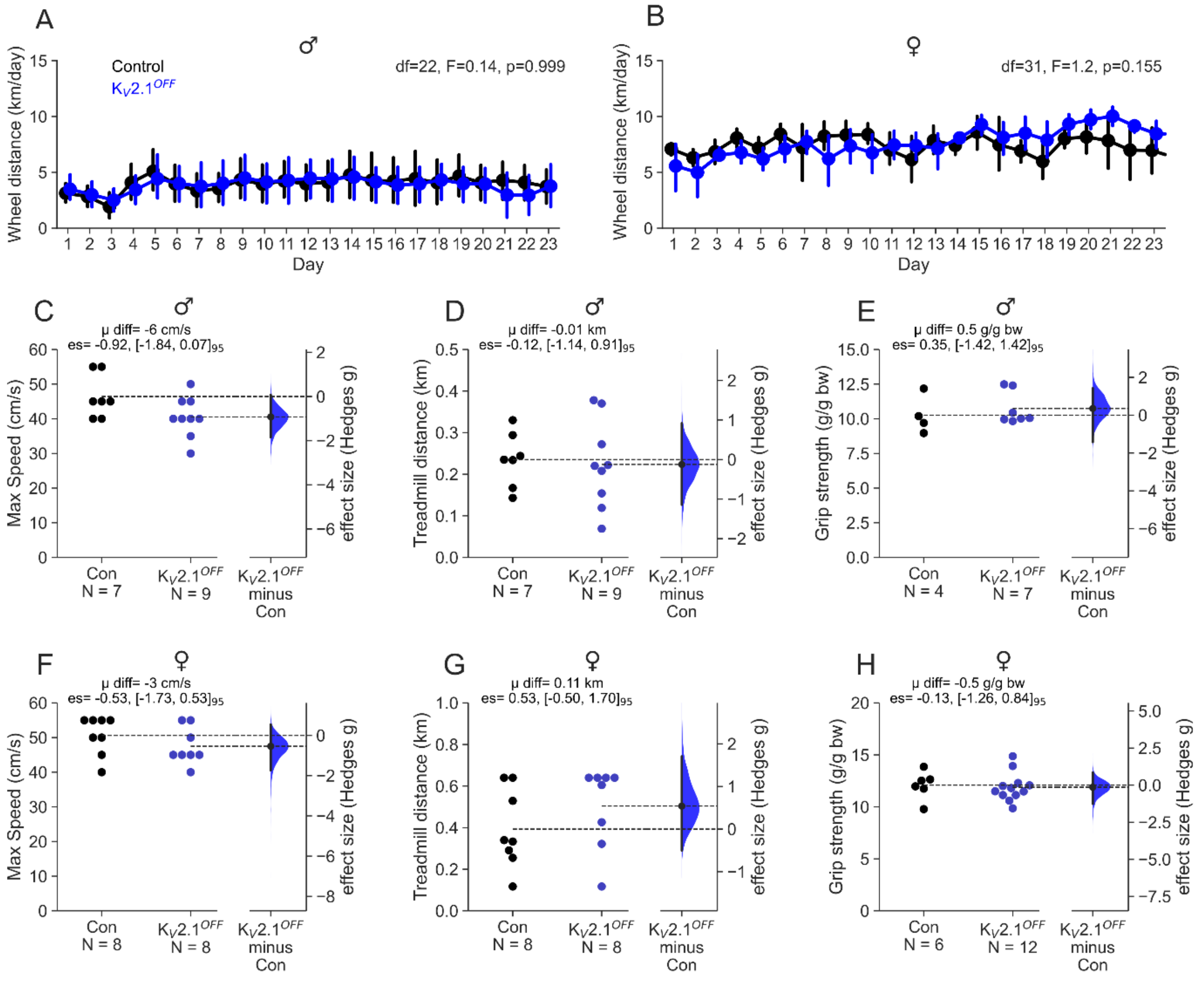


**Figure 9- figure supplement 1. Comparing behaviour in control and ChAT-K$_V$2.1$^{OFF}$ mice .**

**(A-B)** Distance run per day for male **(A)** and female **(B)** control (black; male N=6, female N=5) and ChAT-K$_V$2.1$^{OFF}$ (grey; male N=6, female N=6) mice. Output of repeated measures ANOVA shown at the top of each graph. Maximum speed **(C)** and distance **(D)** run by male mice on a treadmill at 15° incline. **(E)** Grip strength (g) in male mice normalised to body weight (g). **(F-H)** as in **C-E**, but for female mice. Gardner-Altman estimation plots show treadmill and grip strength data in **(C-H)**. Control and ChAT-K$_V$2.1$^{OFF}$ groups are plotted on the left axes and the bootstrapped sampling distribution (5000 reshuffles) for Hedges g effect sizes are plotted on the right. The Hedges g effect size is depicted as a dot; the 95% confidence interval is indicated by the vertical error bar. The mean difference (μ), effect sizes (es) and 95% confidence intervals [lower, upper] are displayed at the top of each plot. Experimental unit (N) = animals.

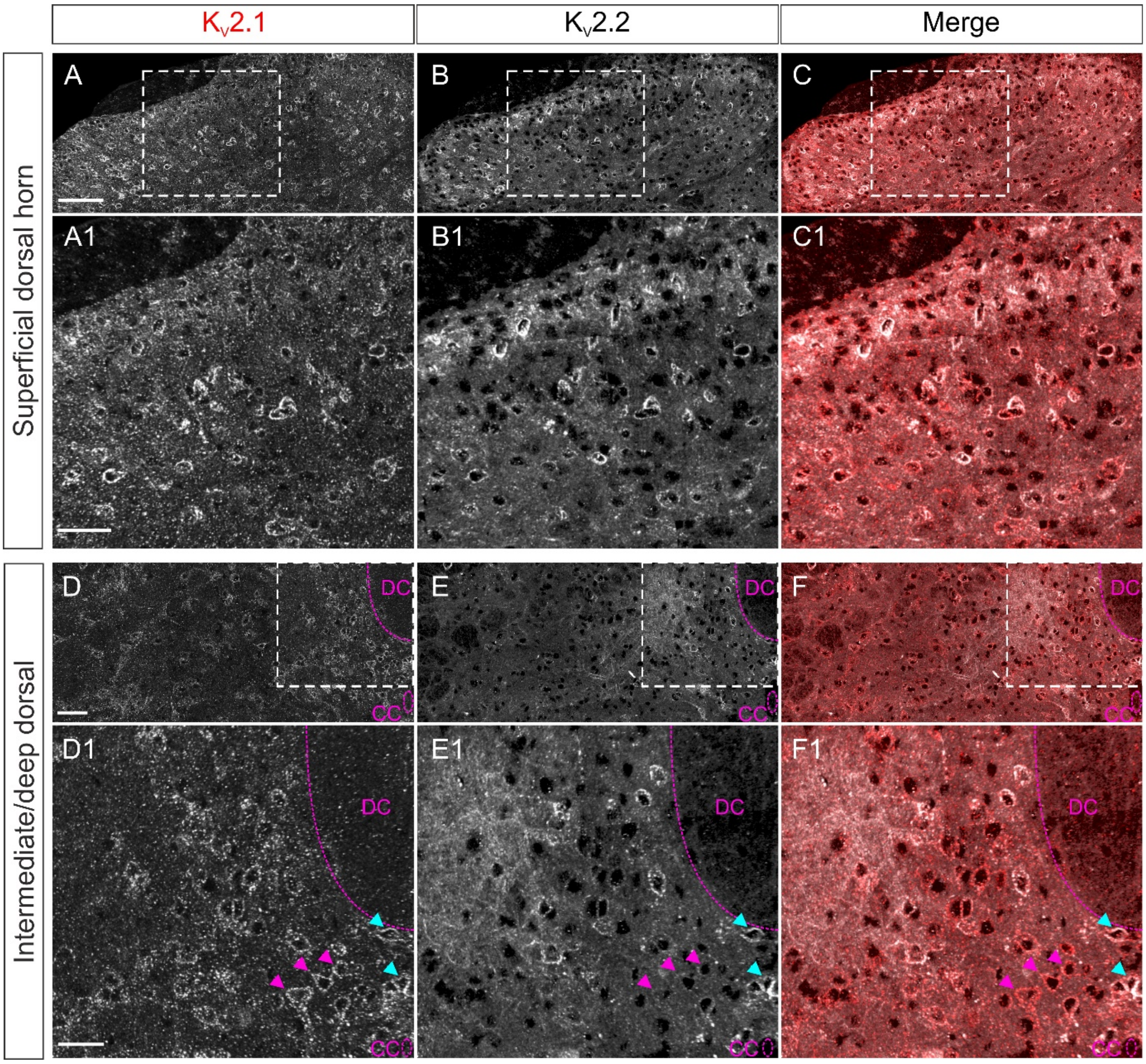


**Figure 10- figure supplement 1. $K_V$2 expression throughout the spinal laminae. (A-F1)** 40x confocal tiled Z stack (5 x 1μm slices) projection images of the superficial laminae of the lumbar spinal cord stained for $K_V$2.1 **(A-A1, D-D1)** and $K_V$2.2 **(B-B1, E-E1)**; merged image shown in **(C-C1, F-F1)**. **(A-C)** Shows the superficial laminae of the dorsal horn, with (A1-C1) showing the region marked by a box in (A-C) cropped and expanded. **(D-F)** Shows the deep dorsal and intermediate laminae, with (D1-F1) showing the region marked by a box in (D-F) cropped and expanded. Magenta arrow heads indicate $K_V$2.1 high, $K_V$2.2 low expression cells. Light blue arrow heads indicate cells expressing high $K_V2.1$ and $K_V2.2$ levels. DC= dorsal columns, CC= central canals. Scale bars in (A-C) = 40 µm, (A1-C1) = 25 µm, (D-F) = 40 µm, (D1-F1) = 20 µm.
